# EegFun.jl: A Julia Package Tutorial for EEG Analysis

**DOI:** 10.64898/2026.08.11.744163

**Authors:** Carolin Dudschig, Samuel Sonntag, Ian G. Mackenzie

## Abstract

`EegFun.jl` is an open-source package for electroencephalography (EEG) analysis implemented in the Julia programming language. `EegFun.jl` provides a flexible framework for EEG research, covering data import from standard file formats, filtering and re-referencing, Independent Component Analysis (ICA) for artifact detection/correction, epoch extraction, and ERP averaging and visualisation. The Julia language provides the readability of a high-level scripting environment together with execution speeds comparable to compiled code. EegFun.jl combines interactive data visualization with high-performance execution, making large-scale analyses both efficient and easy. Here, we provide a brief overview and introductory tutorial of the core stages of the EEG analysis workflow to illustrate the package’s capabilities. The package is freely available under the MIT license.

## Introduction

The electroencephalogram (EEG) records electrical brain activity from electrodes placed on the scalp, capturing the summed post-synaptic potentials of cortical neuron populations. First formally described by Berger (1929), EEG has since become one of the most widely used methods in cognitive neuroscience and clinical practice, valued primarily for its non-invasive nature and its direct, millisecond-resolution measure of neural dynamics (Luck, 2014).

The event-related potential (ERP) technique derives cognitive neural signatures by averaging EEG segments time-locked to stimuli or behavioural events across many trials (Luck, 2014). On the surface, this process is conceptually simple: we continuously record voltage across multiple channels whilst a participant performs a task. Event markers signal points of theoretical interest, around which discrete time-windows or epochs are extracted. These epochs are then averaged for each experimental condition, and specific features of the resulting waveform, known as ERP components (e.g., the P1, N170, or N400), are compared using standard statistics.

However, even for the simplest of ERP studies, such a straightforward description hides a multitude of methodological choices and processing steps. The refined ERP plots presented in final publications are a long way from the raw, noisy continuous EEG data observed during data collection or initial raw-data inspection. Indeed, the ERP itself is entirely invisible in the raw recording. Recovering the signal requires navigating a “garden of forking paths” (Gelman & Loken, 2014): choices regarding reference channels, filter parameters, artifact identification, threshold-based trial rejection, and epoch window boundaries collectively determine the final result and must be explicitly documented for reproducible science (Luck, 2014; Šoškić et al., 2024).

`EegFun.jl` offers comprehensive options to navigate these choices, allowing researchers to piece together highly customised analysis pipelines whilst automatically logging parameters for exact reproducibility. Alongside this flexibility, the package also provides an automated, configuration-driven batch pipeline (see the Automated Batch Pipeline section). Importantly, both approaches operate on the exact same underlying data structures, ensuring that exploratory single-subject analyses can be easily scaled into full-study batch processing.

The field of cognitive neuroscience is already supported by a number of excellent, highly comprehensive open-source analysis toolboxes. MATLAB-based tools such as EEGLAB (Delorme & Makeig, 2004), ERPLAB (Lopez-Calderon & Luck, 2014), and FieldTrip (Oostenveld et al., 2011) have driven decades of research and remain foundational to the field, whilst MNE-Python (Gramfort et al., 2013) provides a robust, modern ecosystem in Python. `EegFun.jl` does not aim to replace these mature ecosystems, but rather to offer an additional, complementary tool. Maintaining interoperability, including support for importing EEGLAB and FieldTrip data structures, allows researchers to integrate Julia’s computational performance directly into their existing workflows.

The development of `EegFun.jl` is driven by three primary motivations. First, whilst the established MATLAB-based toolboxes are themselves open-source, the underlying MATLAB environment requires an expensive proprietary licence. This financial barrier restricts access and hinders computational reproducibility for researchers operating outside of well-funded institutions. Although free alternatives like GNU Octave exist, full compatibility is not always guaranteed. Second, Julia (Bezanson et al., 2017) provides computational scalability that facilitates researchers to transition from users to developers. Whilst excellent Python ecosystems achieve high performance by offloading heavy computations to compiled external libraries (e.g., NumPy, Numba), researchers attempting to implement novel algorithms that cannot be easily expressed through these existing routines often face severe performance bottlenecks in pure Python. By addressing this “two-language problem” by compiling high-level, dynamically typed code directly to efficient machine instructions via LLVM, Julia allows researchers to write custom, efficient algorithms in the same accessible language they use for daily scripting. Third, the package expands the ecosystem of choice, providing researchers with an additional, modern environment tailored for reproducible analysis.

Here we present `EegFun.jl`, an open-source, pure-Julia EEG package. Rather than providing an exhaustive tutorial on EEG methodology, we first describe the package architecture and its core data structures, demonstrate some of its basic functionality, and finally present a complete continuous EEG to ERP preprocessing pipeline on example data, giving readers a practical sense of the package’s design, syntax, and capabilities. Detailed step-by-step tutorials, extended user guides, and a complete API reference are available on the full documentation webpage (https://igmmgi.github.io/EegFun.jl/dev/).

## Software Architecture and Implementation

### Design Philosophy

`EegFun.jl` is organised around three principles. First, *accessibility*: the public API adopts a consistent, function-based style in which each analysis step maps to a single, clearly named function call (e.g., `highpass_filter`, `rereference`, or `extract_epochs`), keeping the analysis logic readable even for researchers with limited programming experience. Second, *transparency*: every processing step is implemented in pure Julia, meaning the full source code is available for inspection, modification, and extension without leaving the host language. Third, *reproducibility*: by encouraging scripted rather than GUI-driven workflows, `EegFun.jl` ensures that a complete analysis can be re-executed exactly from a single script file or configu, satisfying open-science requirements for computational reproducibility.

Additionally, `EegFun.jl` was designed to be highly interactive, making it easy to visually inspect data without leaving the Julia environment. Where GUIs are provided, they complement this exploratory process rather than replace the scripting API, ensuring that every GUI action has a direct scripted equivalent.

### The Julia Language

Researchers transitioning to Julia will interact primarily with the Julia Read-Eval-Print Loop (REPL), or via interactive notebooks (such as Jupyter or Pluto) and integrated development environments (like VS Code). Like Python or MATLAB, Julia code can be evaluated interactively line-by-line, allowing for rapid, exploratory data analysis. Basic operations use a familiar `function_name(data, parameters)` syntax. Because of these structural similarities, researchers familiar with Python or MATLAB should find the transition to Julia straightforward.

Beyond core syntax, a major advantage of the Julia ecosystem for open science is its built-in package manager (`Pkg`), accessible directly from the REPL by pressing the ] key. By automatically recording the exact versions of all installed dependencies in `Project.toml` and `Manifest.toml` files, Julia ensures that an analysis environment constructed today can be replicated by other researchers in the future, eliminating the dependency-conflict issues common in other scripting ecosystems. Additionally, the REPL provides a dedicated help mode (accessed by pressing `?`), allowing researchers to pull up detailed documentation for any function without leaving their analysis environment. Despite these underlying features, `EegFun.jl` remains highly accessible; for users without prior programming experience, a complete analysis can be executed with very little actual programming beyond typing a sequence of straightforward commands.

### Software Dependencies

`EegFun.jl` operates entirely within the native Julia environment and does not require a MATLAB or Python installation. Under the hood, `EegFun.jl` utilises several core Julia packages. For visualisation, *Makie.jl* (Danisch & Krumbiegel, 2021) provides an interactive and publication-quality plotting layer through its GLMakie (GPU-accelerated) and CairoMakie (vector-graphics) backends. Core signal processing capabilities, including FIR/IIR filter design and fast Fourier transforms, are powered by *DSP.jl* and *FFTW.jl*, respectively. For data management, *DataFrames.jl* (Bouchet-Valat & Kamiński, 2023) serves as the core backbone of the package for data organisation, storing all EEG time-series data and metadata in intuitive, easy-to-follow tabular formats, whilst *JLD2.jl* handles fast, HDF5-compatible serialisation to disk.

### Supported File Formats

`EegFun.jl` can import data from common EEG file formats including Biosemi Data Format (`.bdf`), the European Data Format (`.edf`), the Extensible Data Format (`.xdf`), and the Functional Image Format (`.fif`). To ensure interoperability with existing analysis ecosystems, it also includes readers for EEGLAB `.set` files (Delorme & Makeig, 2004), the BrainVision `.vhdr` format, and FieldTrip `.mat` structures (Oostenveld et al., 2011) (the latter supported via the *MAT.jl* package). Processed data structures can be quickly saved to disk and reloaded using the HDF5-compatible `.jld2` format. Additionally, the package supports exporting datasets directly into Brain Imaging Data Structure (BIDS) compliant folder architectures to facilitate open-science sharing.

### Core Data Structures

The package defines a formal, typed hierarchy of data structures to represent EEG data at successive stages of the analysis pipeline (Table 2). In Julia, these custom types are implemented as `structs`, which are composite containers defined within the package that group related variables (such as time-series matrices and spatial metadata) into a single, cohesive data structure. The workflow typically begins with the `RawData` type, which holds the unprocessed, continuous recording as read from disk, including the raw sample matrix and header metadata. This is subsequently transformed into `ContinuousData`, an annotated struct that explicitly binds the time-series matrix with a *Layout* structure containing 2D and 3D channel coordinates, channel labels, and event annotations. Channel layout files (`.csv`) are read separately and combined with the EEG data at this construction stage, enforcing that critical spatial metadata is never silently discarded. Following segmentation around event triggers or markers, the data becomes `EpochData`, preserving individual trials alongside condition labels and per-trial metadata. Averaging these trials yields `ErpData`. The package also defines structures for more advanced analyses, such as `TimeFreqData` and `TimeFreqEpochData`, which store multi-dimensional spectral power and phase information.

At its core, the time-series data within each of these structures is stored as a `DataFrames.jl DataFrame` (Bouchet-Valat & Kamiński, 2023) in which rows correspond to time points and columns correspond to named channels. Because this underlying structure is simply a standard `DataFrame` (Table 1), it offers extensive flexibility; researchers can easily store additional arbitrary data, such as trial-level reaction times or artifact masks, under new column headers. An explicit `time` column records the latency of every sample in seconds, so the temporal axis is always unambiguous and never needs to be reconstructed from sample rate and offset. This tabular layout keeps the data model deliberately simple: channels are accessed by their symbolic labels (e.g., `:Cz`, `:Pz`) rather than by numeric indices, eliminating a common source of off-by-one and transposition errors that can occur when working with raw channels *×* timepoints matrices. Standard `DataFrame` operations, including column selection, row filtering, joins, and CSV export, work directly on the EEG data, making it straightforward to interface with external statistical packages or to inspect the data interactively in the Julia REPL. For segmented data, `EpochData` extends this pattern by storing each trial as a separate `DataFrame` in a `Vector`, keeping the per-trial structure explicit and making trial-level operations (e.g., rejection, reordering, or condition subsetting) naturally expressible as standard vector operations. To facilitate this, `EegFun.jl` provides dedicated built-in functionality that makes executing these common trial-level manipulations straightforward.

**Table 1.** The internal DataFrame representation of continuous EEG data, demonstrating the explicit time and sample channels alongside channel voltage columns.

| time | sample | trigger | Fp1 | AF7 | AF3 | F1 | ... |
| --- | --- | --- | --- | --- | --- | --- | --- |
| 0.0000 | 1 | 0 | -1544.81 | 5186.93 | -5627.15 | 873.31 | ... |
| 0.0019 | 2 | 0 | -1548.72 | 5180.24 | -5630.05 | 871.44 | ... |
| ⋮ | ⋮ | ⋮ | ⋮ | ⋮ | ⋮ | ⋮ | ⋮ |

**Table 2.** Primary Data Structures in EegFun.jl.

| Concrete Type | Abstract Type | Description |
| --- | --- | --- |
| RawData | — | Unprocessed raw data as read from disk. |
| ContinuousData | SingleDataFrameEeg | Continuous recording with layout and metadata. |
| EpochData | MultiDataFrameEeg | Segmented trial data (vector of dataframes). |
| ErpData | SingleDataFrameEeg | Event-related potentials averaged across trials. |
| TimeFreqData | EegFunData | Averaged time-frequency power (and phase). |
| TimeFreqEpochData | EegFunData | Single-trial time-frequency decompositions. |
| AnalysisInfo | — | Analysis metadata. |

The function `create_eegfun_data` enforces that the time-series data, the spatial *Layout*, and an `AnalysisInfo` metadata record are always bundled together (Listing 1^1^). This prevents the silent loss of channel coordinates that commonly occurs when data are passed between separate variables in other toolboxes. As a result, spatially demanding operations, such as spherical spline interpolation for bad-channel reconstruction, always have the metadata they require. This typed hierarchy also enables `EegFun.jl` to exploit Julia’s *multiple dispatch* mechanism, in which a single function name automatically selects the correct implementation based on the run-time types of its arguments. For example, the `baseline` function can be called identically on an `EpochData` instance (correcting each trial individually), on an `ErpData` instance (correcting the averaged waveform), or on a `Vector{EpochData}` (iterating over multiple conditions). In each case the user writes the same call, such as `baseline(dat, (−0.2, 0.0))`, and Julia dispatches to a specialised method that handles the specific type correctly. This dispatch pattern is applied consistently throughout the entire package. By relying on multiple dispatch rather than complex runtime `if`-`else` branches, the code remains both extensible and efficient: adding support for a new data type requires only a new method definition, without modifying any existing code.

### Data Selection and Predicates

A core design principle of `EegFun.jl` is the strict semantic separation between temporal windowing and spatial-level subsetting. Whilst temporal calculation windows (such as baseline correction intervals) are defined using native Julia tuples representing time in seconds (e.g., `(−0.2, 0.0)`), this subsetting is performed via a dedicated predicate-based selection API. Functions throughout the package accept `channel_selection` and `sample_selection` arguments to subset data for specific operations, metrics, or visualisations.

Instead of relying on hardcoded numeric indices, which are fragile and prone to silent errors when processing layouts with different channel counts, selections are created using dedicated helper functions like `EegFun.channels()` and `EegFun.samples()`. These functions return logical predicates that are subsequently applied to the underlying tabular data structure. For example, a researcher can restrict an analysis to specific channels by passing an array of symbolic channel labels (e.g., `channel_selection = EegFun.channels([:Fz, :Cz, :Pz])`), filter samples based on precomputed metadata flags (e.g., `sample_selection = EegFun.samples(:is_artifact_free)`), or invert selections to exclude specific data points (e.g., `EegFun.samples_not(:extreme)`). These helper functions accept custom user-defined functions as arguments, enabling dynamic, programmatic subsetting (e.g., passing a string-matching function to dynamically select all left-hemisphere channels, as demonstrated in Listing 6). This approach ensures that subsetting logic remains readable, reproducible, and integrated with Julia’s multiple dispatch system, allowing the same selection syntax to be used consistently across continuous, epoched, and averaged data structures. Because selections are first-class objects rather than ad-hoc index vectors, they can be defined once and reused across every operation in an analysis script, guaranteeing consistency and reducing the opportunity for copy-paste errors (Listing 2).

**Listing 1:**
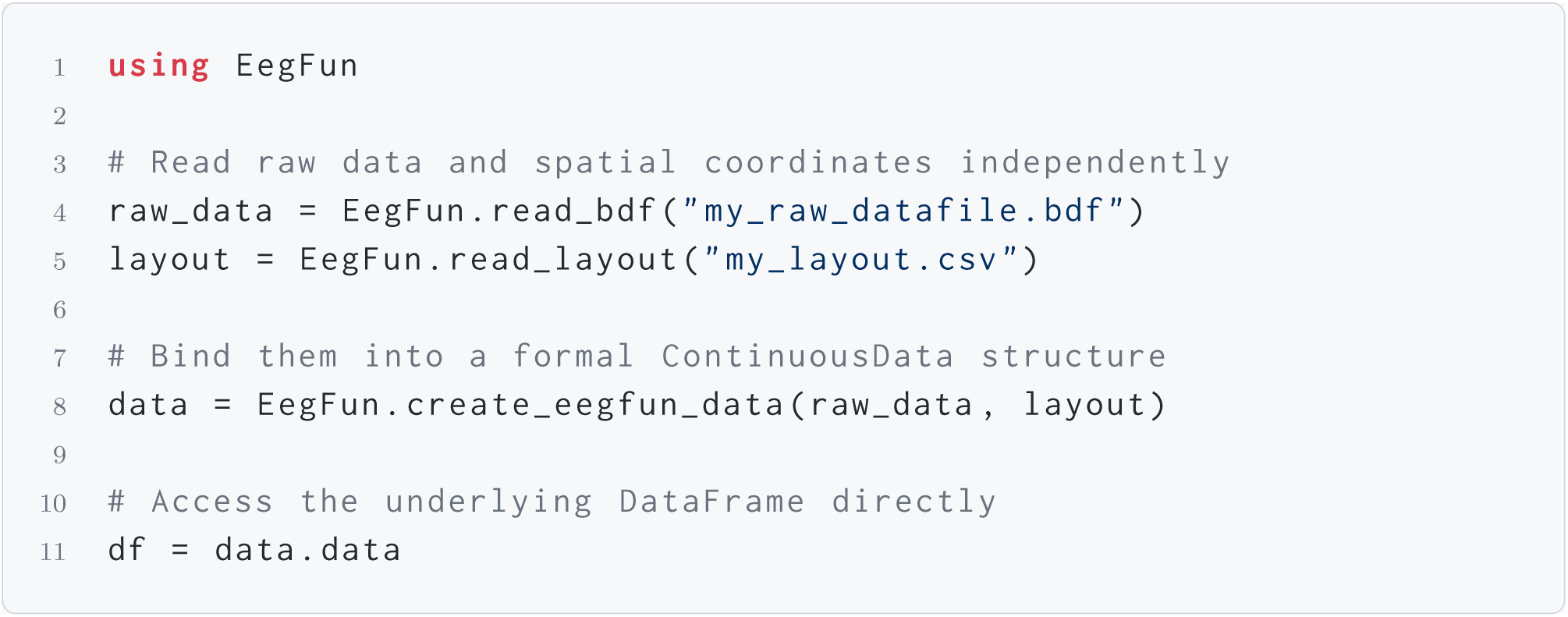
Constructing a `ContinuousData` instance from raw data and a layout file.

### Core Functionality

`EegFun.jl` covers the full EEG to ERP analysis workflow through several tightly integrated functional areas. For data I/O, format-specific readers follow a unified `read_*` prefix convention alongside robust persistence mechanisms to and from `.jld2` files. The preprocessing suite provides a comprehensive set of signal processing routines, from standard filtering and rereferencing to robust artifact management. Data cleaning is supported by both automated sensor-space thresholding and Infomax ICA decomposition (Ablin et al., 2018; Bell & Sejnowski, 1995; Jung et al., 1998), complete with tools for automated EOG identification and interactive review. The core epoching and averaging routines provide flexible trigger-sequence matching through `EpochCondition` configurations, baseline correction, absolute threshold and statistical-based trial rejection, and standard arithmetic ERP averaging. These operations are complemented by an extensive visualisation system containing an interactive continuous data browser, ERP waveform and topographic plots, an ICA component viewer, and a combined ERP–topography layout display. Beyond these core operations, the package also supports advanced analyses including Morlet wavelet, multitaper, and STFT time-frequency decompositions, as well as multivariate pattern analysis (MVPA) with cross-validated decoding.

**Listing 2:**
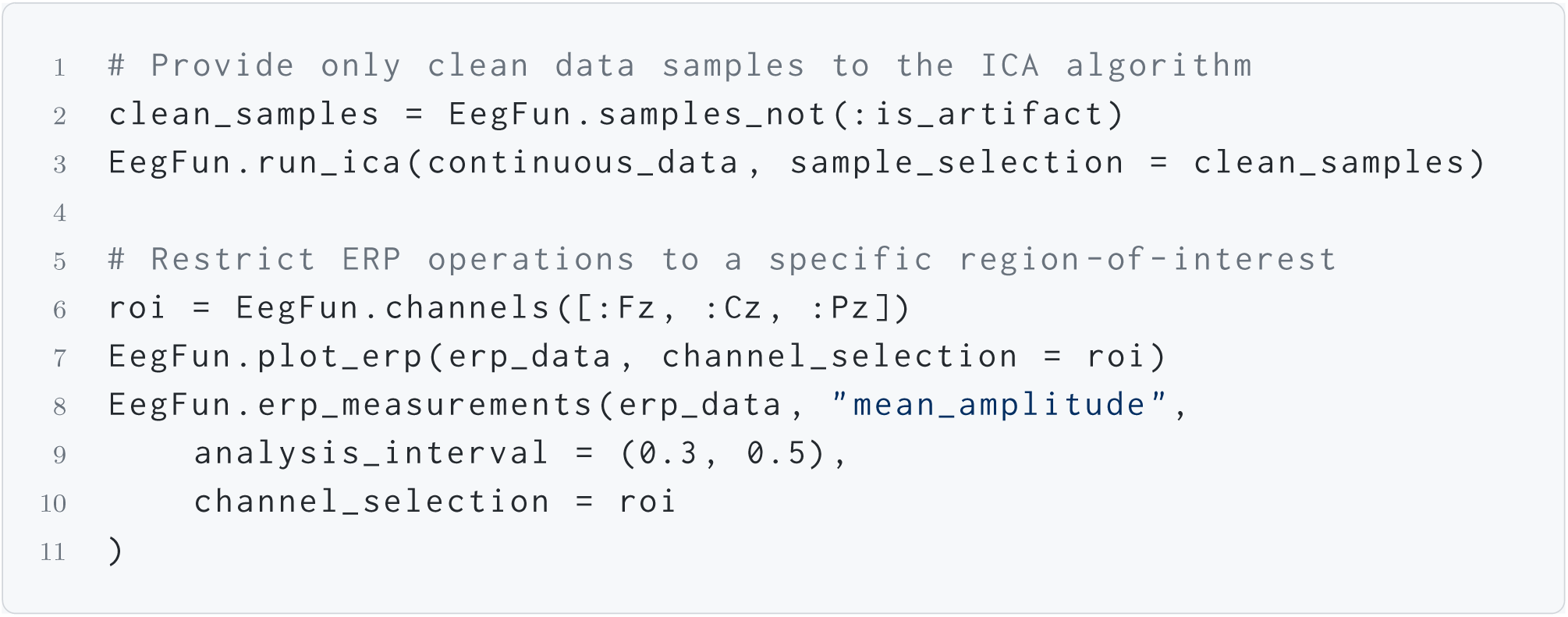
Predicate-based data selection: composable, self-documenting, and reusable across all package operations.

### Availability

`EegFun.jl` is freely available under the MIT licence on GitHub (https://github.com/igmmgi/EegFun.jl). The package is registered in the Julia General Registry and can be installed in a single command:

`using Pkg; Pkg. add(” EegFun “)`

Full online documentation, including a complete API reference and a suite of worked examples covering the preprocessing pipeline, time-frequency analysis, and MVPA decoding, is available at https://igmmgi.github.io/EegFun.jl/dev/. The documentation features dedicated preprocessing guides and complete experiment walkthroughs that take users from raw data through to statistical inference. All example datasets and layout files required to reproduce these tutorials are openly hosted on Zenodo (https://doi.org/10.5281/zenodo.21770157). Rather than requiring manual downloads, `EegFun.jl` leverages the `DataDeps.jl` framework to automatically download and extract these datasets the first time a user requests them.

## Example Workflows

To accommodate both exploratory analysis and high-throughput execution, `EegFun.jl` supports two distinct workflow paradigms. The following sections first demonstrate an interactive, step-by-step approach for single-subject analysis, followed by an automated, configuration-driven batch pipeline for full studies. Importantly, these examples are intended as a concise feature overview rather than a comprehensive tutorial; they highlight representative capabilities of the package without covering every available function or parameter. More detailed tutorials, including instructions on how to use the built-in `DataDeps.jl` integration to automatically download sample datasets, are available in the online documentation at https://igmmgi.github.io/EegFun.jl/dev/.

### Interactive Step-by-Step Walkthrough

This approach allows researchers to dynamically piece together individual analysis components. It is suited for data exploration, parameter tuning, and teaching, as it provides students and researchers new to EEG analysis with a transparent view of the underlying signal processing steps and methodological decision points. After successfully installing the Julia language and the EegFun package (see above), the following section provides a step-by-step introduction to a basic EEG analysis, followed by an overview of the full capabilities of the EegFun package.

#### Data Import and Spatial Layout

A critical initial step in the analysis pipeline is associating the raw time-series data with a validated spatial layout. To this end, `EegFun.jl` contains many predefined layouts that can be easily loaded (see https://igmmgi.github.io/EegFun.jl/dev/explanations/layouts). These explicit layouts are required for topographic plots and for defining neighbouring channels (Listing 3). The resulting `ContinuousData` instance bundles the time-series array, the explicit spatial layout, sampling rate, and event annotations. The interactive data browser (Figure 2) can then be launched immediately to inspect the recording. Alternatively, for users preferring a zero-code entry point, `EegFun.jl` provides a unified graphical user interface (`EegFun.plot_gui()`, Figure 3). This GUI interface researchers to interactively load processed `.jld2` or raw data files, visually select channel configurations, and launch any of the package’s plotting functions (such as ERP topographies, ICA component viewers, or the raw data browser) without writing a single line of script.

**Figure 1.**
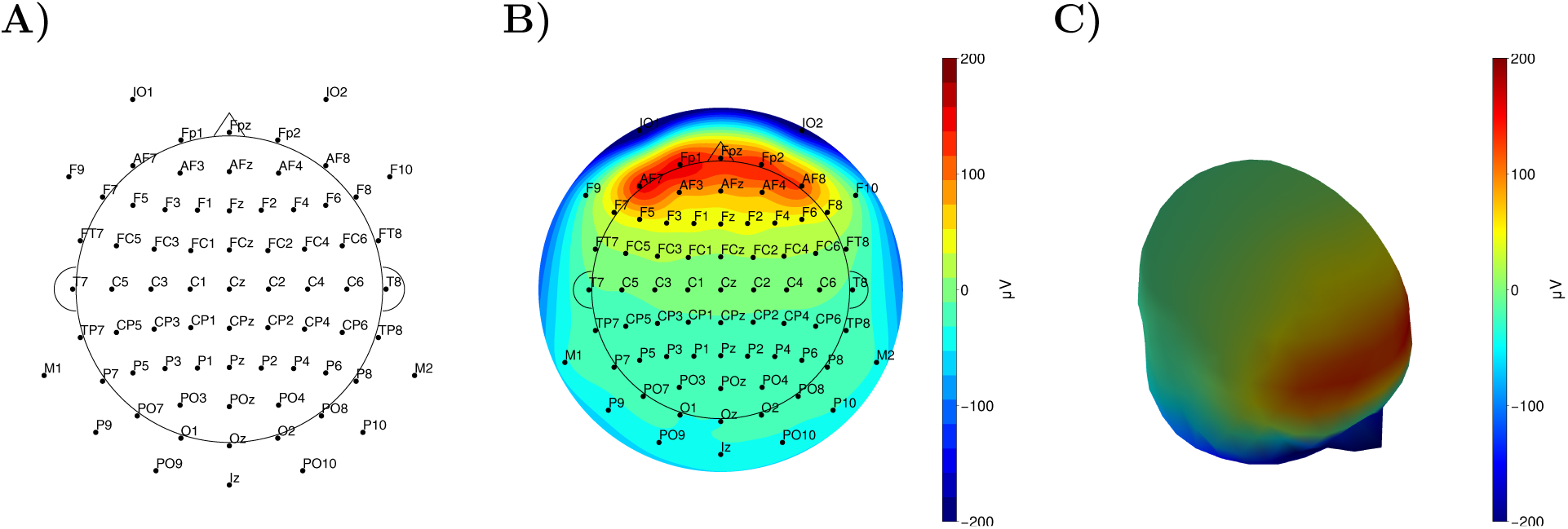
**(A)** 2D spatial mapping of the loaded channel layout. **(B)** 2D topographical interpolation of EEG voltage over a flattened scalp projection. **(C)** 3D topographical interpolation of EEG voltage over a spherical scalp model. The code used to generate these plots is shown in Listing 3.

**Figure 2.**
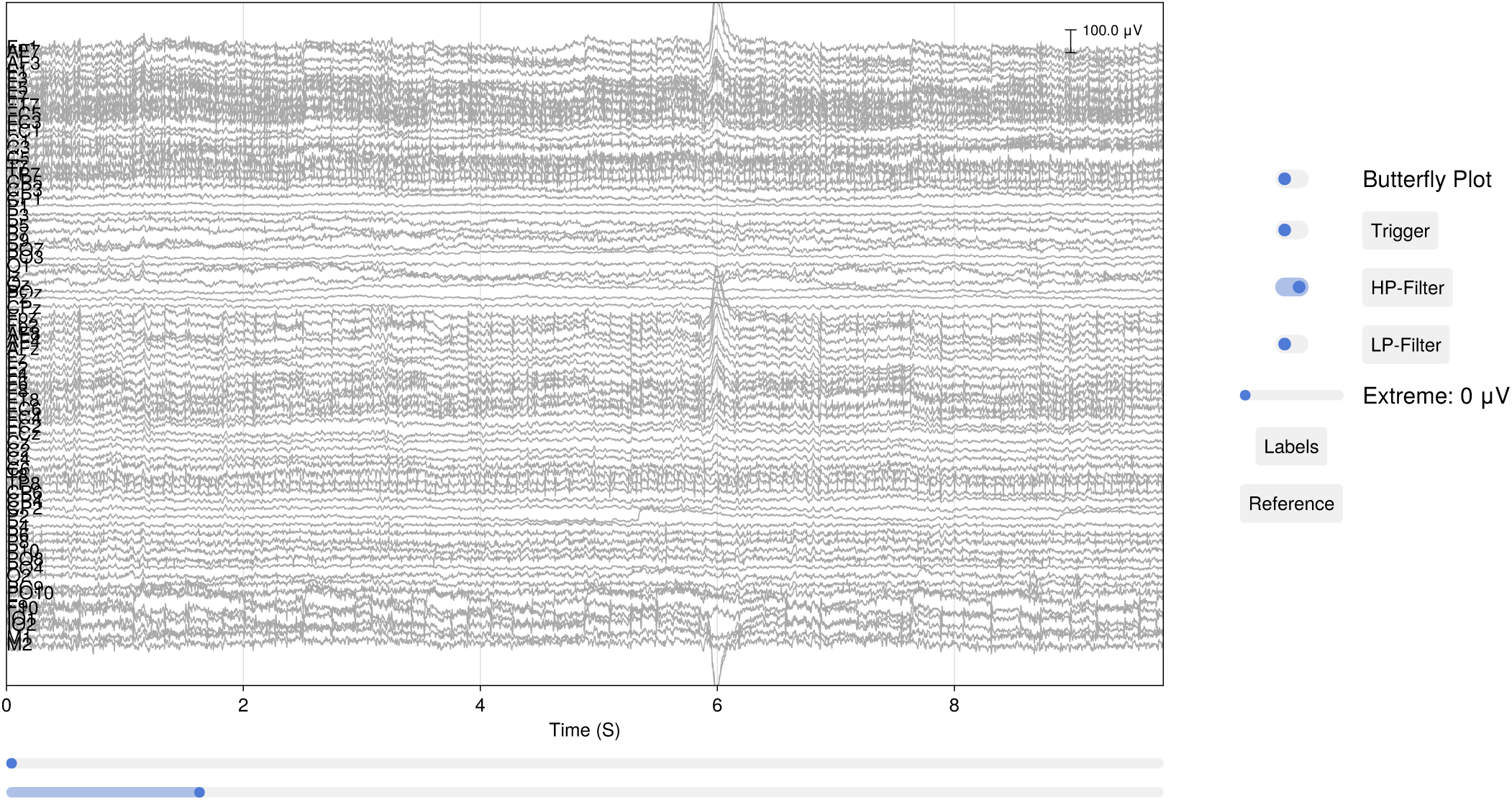
Interactive data browser displaying continuous EEG data. Researchers can scroll through time, adjust the displayed time window, highlight extreme values, toggle or configure online filters, and dynamically change the reference channel. Additional non-EEG channels can be viewed simultaneously and are plotted according to their data type. Specific temporal regions can also be selected directly on the trace using keyboard and mouse combinations. Pressing the `I` key opens a help menu in the REPL detailing all available key combinations, a feature common across several of `EegFun.jl`’s highly interactive plots.

**Figure 3.**
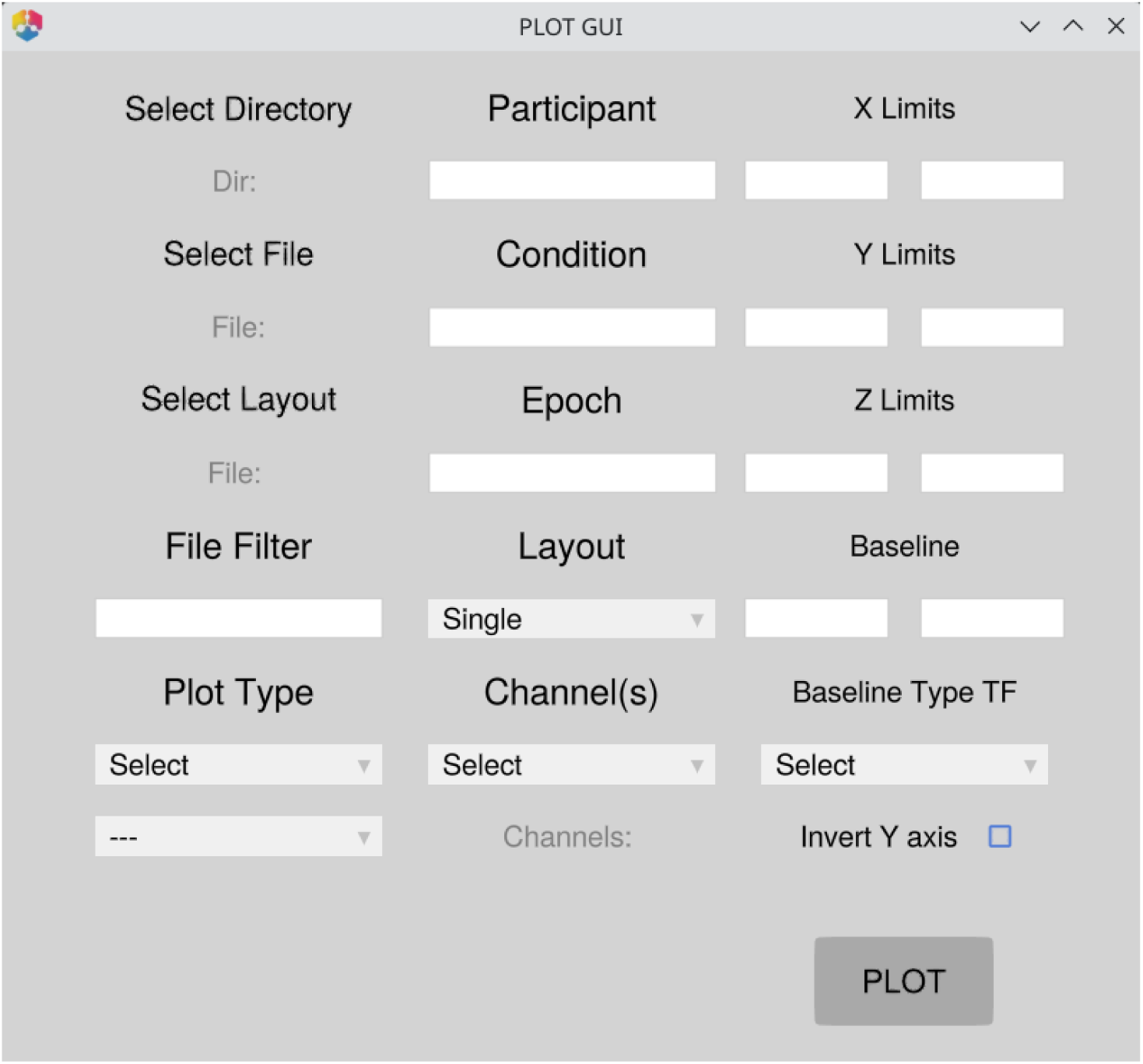
The `EegFun.jl` master GUI provides a zero-code entry point for loading data, configuring analysis parameters, and launching any of the package’s interactive plotting functions.

#### Rereferencing and Filtering

Once the raw data has been imported and the appropriate layout loaded, it is important to visually inspect the data in the time domain (Figure 2) and the frequency domain (Figure 4). Following this inspection, the next stage of the interactive pipeline involves initial preprocessing steps, including re-referencing, and high-pass filtering to remove slow voltage drifts.

**Figure 4.**
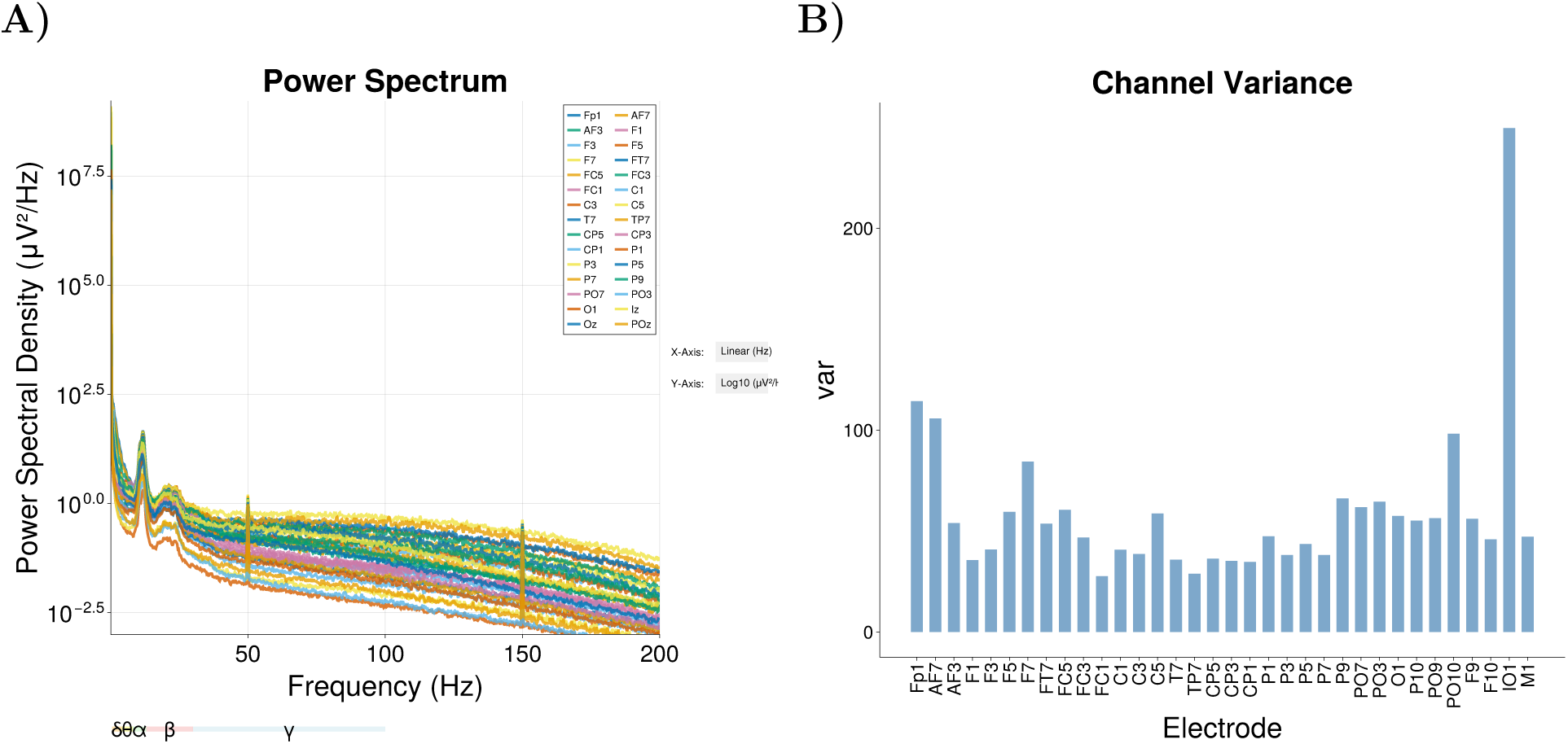
Interactive diagnostic plots for channel-level inspection. **(A)** Channel summary showing the frequency spectrum of the continuous recording. Peaks at 50 Hz line noise frequencies can be easily identified. **(B)** Overview of channel statistics (e.g., variance) providing a complementary method for identifying excessively noisy or flat channels. Here we can see channel IO1 (below left eye) showing higher variance, likely due to eye-blink activity.

For re-referencing, it is important to note that EEG voltages are inherently measured relative to a chosen reference channel. Different recording systems apply different online referencing settings (e.g., the vertex or a mastoid channel) that bias the spatial distribution of recorded activity. However, because re-referencing is a linear operation, researchers can always convert back and forth between different references offline. A common correction is to re-reference to the common average of all scalp channels (average reference), which is often the standard choice for high-density arrays. Alternatively, referencing to linked mastoids or a single channel (e.g., Cz) is also common depending on the research question (see Listing 4 for examples).

**Listing 3:**
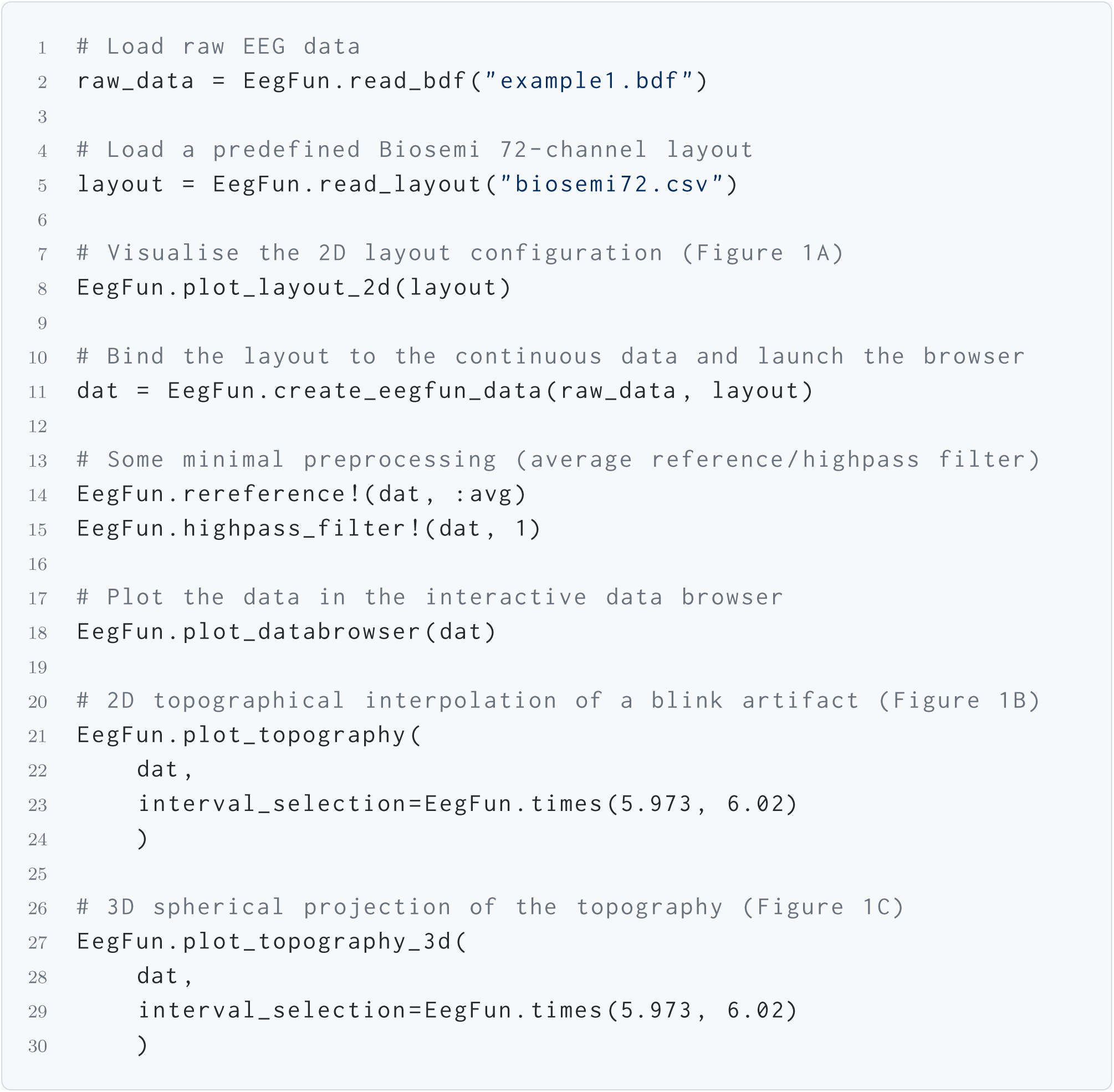
Loading raw data alongside an explicit spatial layout to generate 2D and 3D topographies (Figure 1A–C), and launching the interactive data browser (Figure 2).

For high-pass filtering, slow voltage drifts caused by skin sweating, amplifier DC offsets, or cable movement must be removed, as they contaminate the EEG baseline. A common recommendation for continuous EEG data is to apply a high-pass filter with a cutoff between 0.01 Hz and 0.1 Hz (Listing 5)^2^.

To assist researchers and students in understanding the impact of these choices, `EegFun.jl` includes an interactive filtering dashboard (`EegFun.plot_erp_filter_gui`). This interface provides a clear, hands-on demonstration of how different parameters (such as Butterworth vs. FIR methods, cutoff frequencies, and filter orders) affect the resulting time-domain waveform, allowing users to visually validate their preprocessing strategy before applying it to an entire dataset.

**Listing 4:**
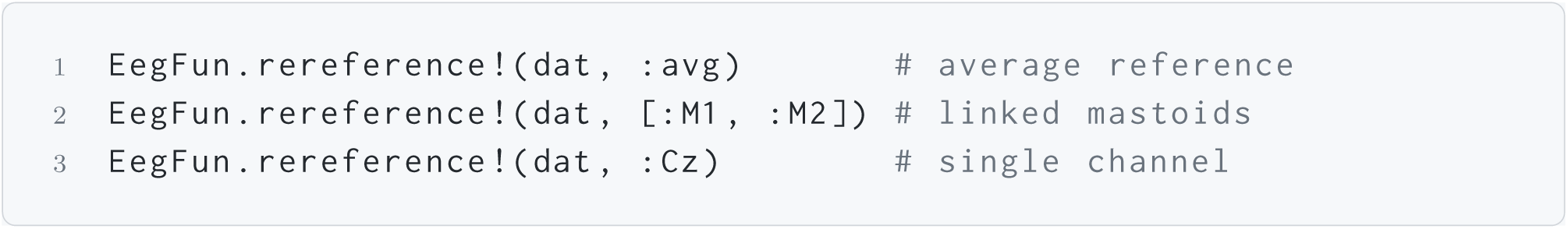
Examples of continuous EEG rereferencing: common average, linked mastoids, and a single channel.

**Listing 5:**
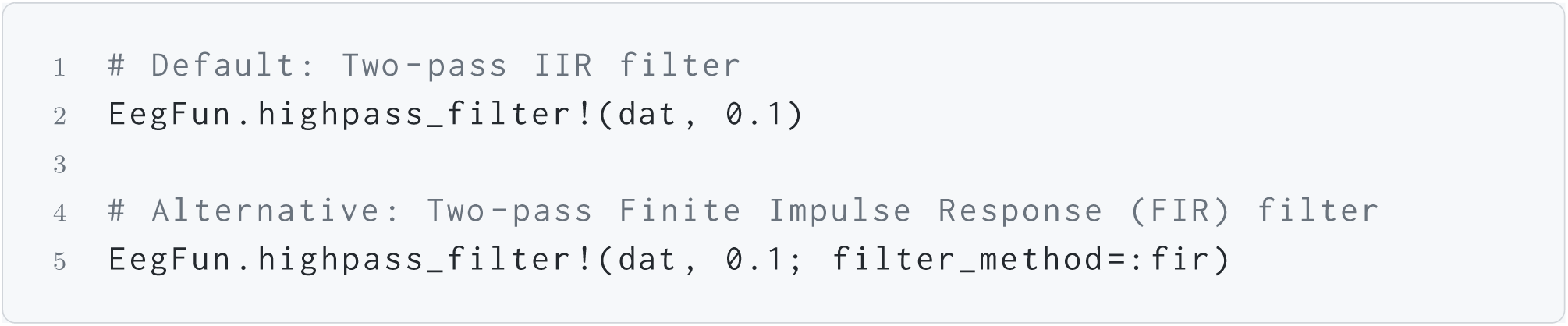
Applying a high-pass filter. By default, the filter applied is a two-pass zero-phase Infinite Impulse Response (IIR) filter via `DSP.filtfilt`, but other options are available.

A visual summary of the channel frequency spectra and basic signal statistics (Figure 4) allows researchers to quickly diagnose line noise or broadband contamination, as well as identify flat or excessively noisy channels prior to downstream analyses (Listing 6).

#### Artifact Correction via ICA

EEG data are prone to contamination from ocular, muscular, and cardiac sources. Rather than rejecting entire data segments, Independent Component Analysis (ICA) decomposes the multichannel signal into statistically independent spatial sources. This allows researchers to selectively remove only those components judged to be artifactual in nature, preserving the remaining neural data for analysis.

ICA requires that source topographies be stationary over time. Low-frequency drift violates this assumption and leads to “ghost components” that absorb slow oscillatory content instead of isolating true artifact sources. A solution is to train ICA on data to which a stronger high-pass filter (e.g., 1 Hz) has been applied (Winkler et al., 2015), and then apply the derived spatial weights back to the original 0.1 Hz recording (Listing 7). The derived Independent Components and their scalp topographies can be inspected interactively (Figure 5A). Furthermore, researchers can examine the continuous activation traces of these components (Figure 5B) and directly overlay the projected component activity onto the raw EEG data browser to verify artifact removal visually (Figure 6). Once the components have been verified visually, researchers can use the automated identification functionality to programmatically flag components matching known artifact signatures (e.g., high temporal correlation with EOG channels or spectral peaks at 50 Hz line noise frequencies). These flagged components can then be removed from the continuous dataset (Listing 8).

**Figure 5.**
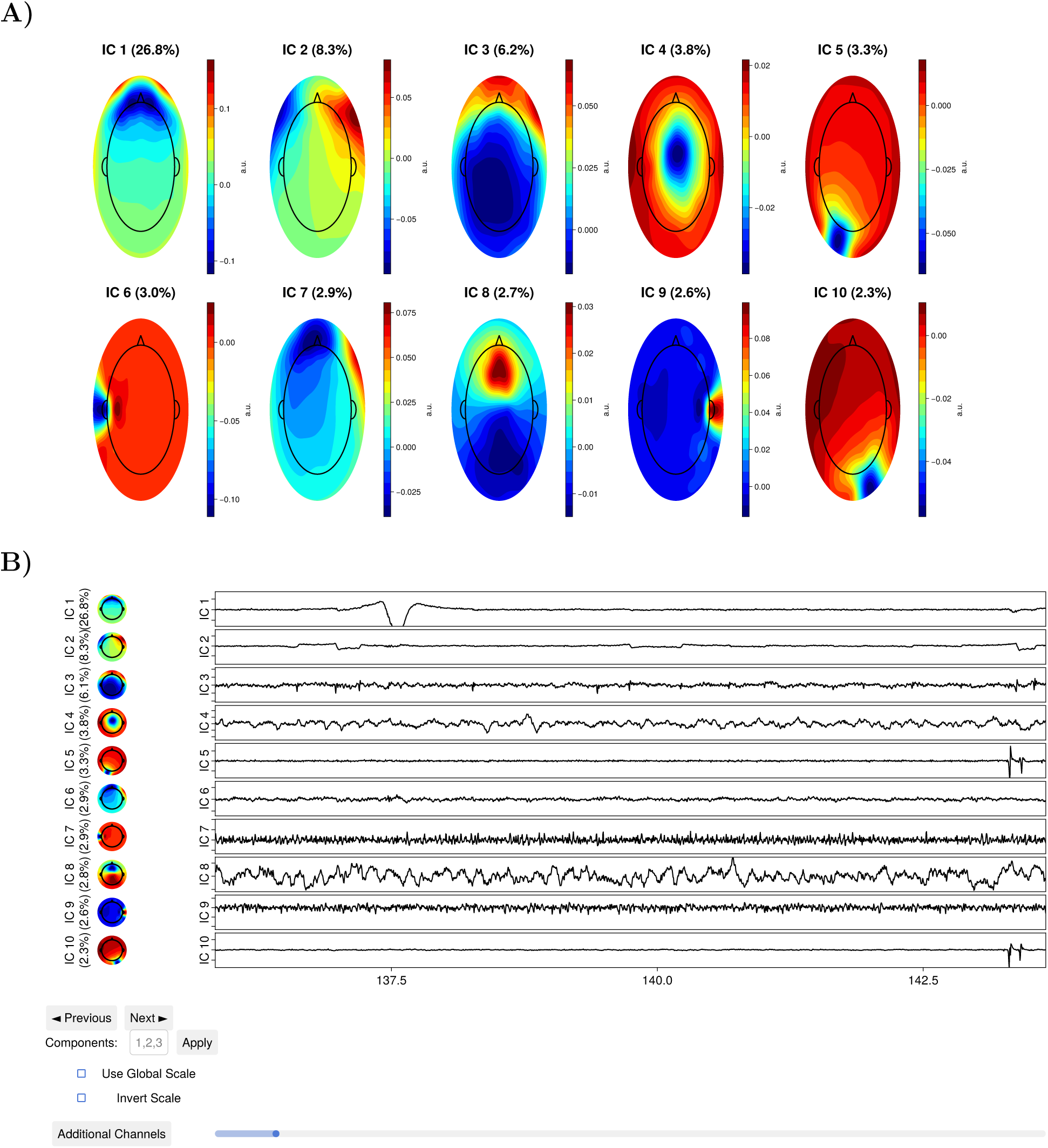
Interactive Independent Component Analysis (ICA) diagnostic views. **(A)** Topographical projections of Independent Components used to identify spatial artifact distributions. For example, eye blinks typically manifest as a strong, focal distribution over the frontal poles (e.g., IC1). **(B)** Continuous activation traces for evaluating temporal dynamics of isolated components.

**Figure 6.**
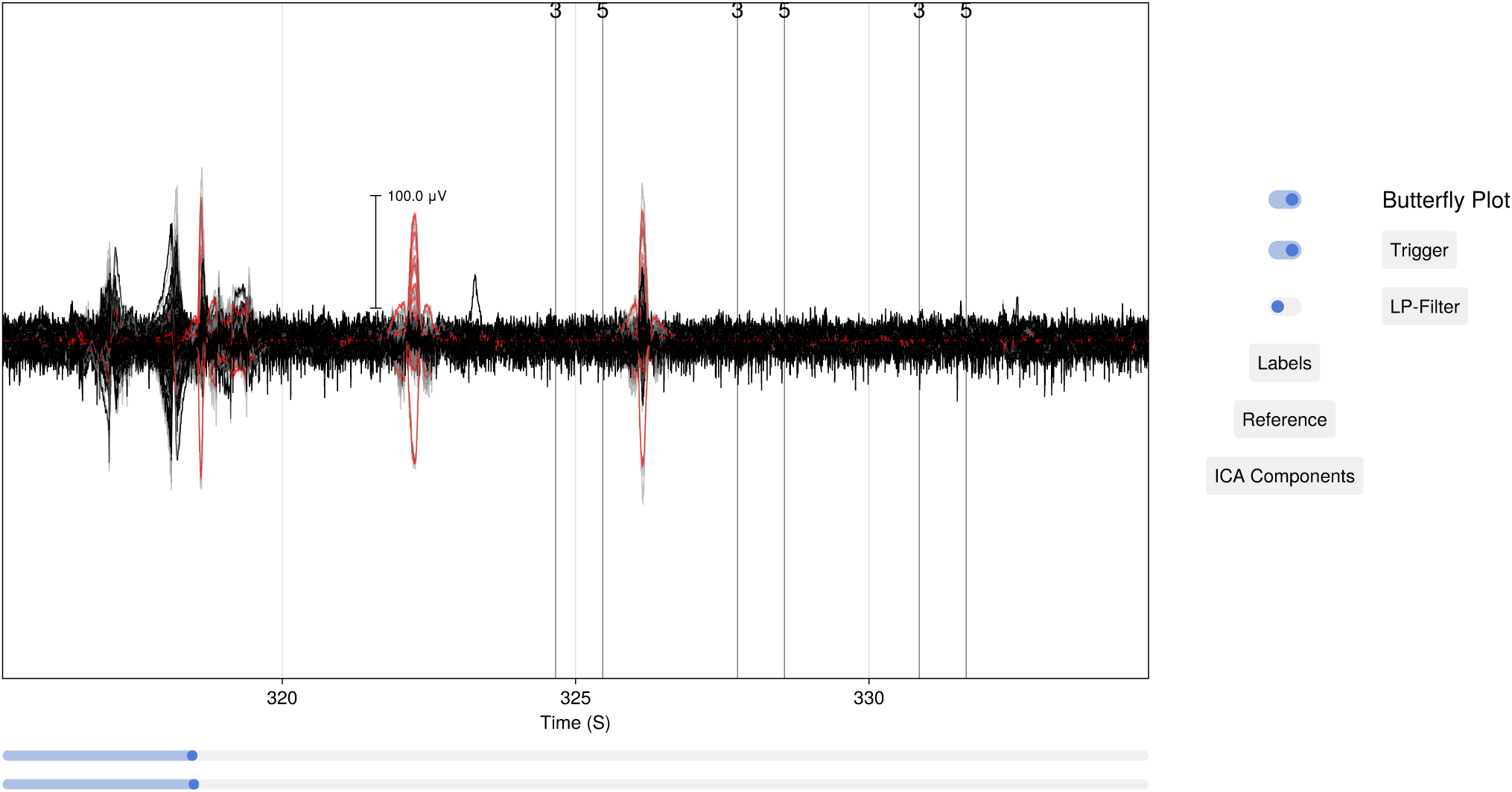
The interactive data browser overlaying the raw EEG data with projected ICA component activity. This view allows researchers to toggle individual artifact components on or off to evaluate their contribution. By utilising the display options, researchers can simultaneously visualise the original, uncorrected EEG trace (plotted in grey), the continuous activation of the selected artifactual components (the subtraction, plotted in red), and the resulting “cleaned” signal (plotted in black).

**Listing 6:**
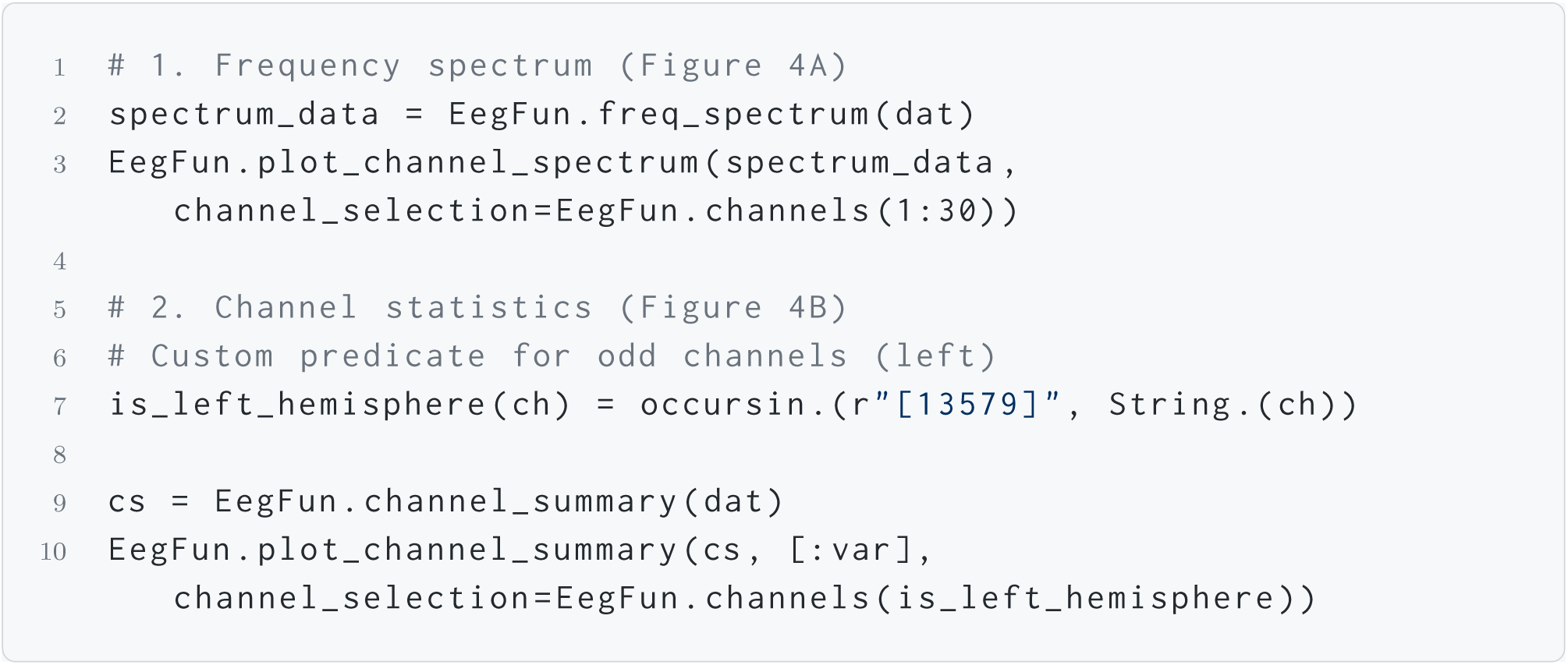
Generating channel frequency spectra and summary statistics to diagnose noise (Figure 4).

#### Epoching, Baseline Correction, and Trial Rejection

Event-Related Potentials (ERPs) are derived by averaging multiple segments of EEG data that are time-locked to specific experimental events. During continuous recording, hardware markers (triggers) are injected into the dataset to precisely document the onset of stimuli or participant responses. To isolate the neural activity associated with these events, researchers extract discrete intervals (epochs) of data around each trigger, typically encompassing a short pre-stimulus period and a longer post-stimulus window (e.g., −500 ms to 1500 ms). `EegFun.jl` offers flexibility in this domain, allowing researchers to define epoch extraction not just based on single triggers, but on sequential trigger patterns, logical combinations, or by applying reaction-time (RT) filtering criteria directly during selection.

**Listing 7:**
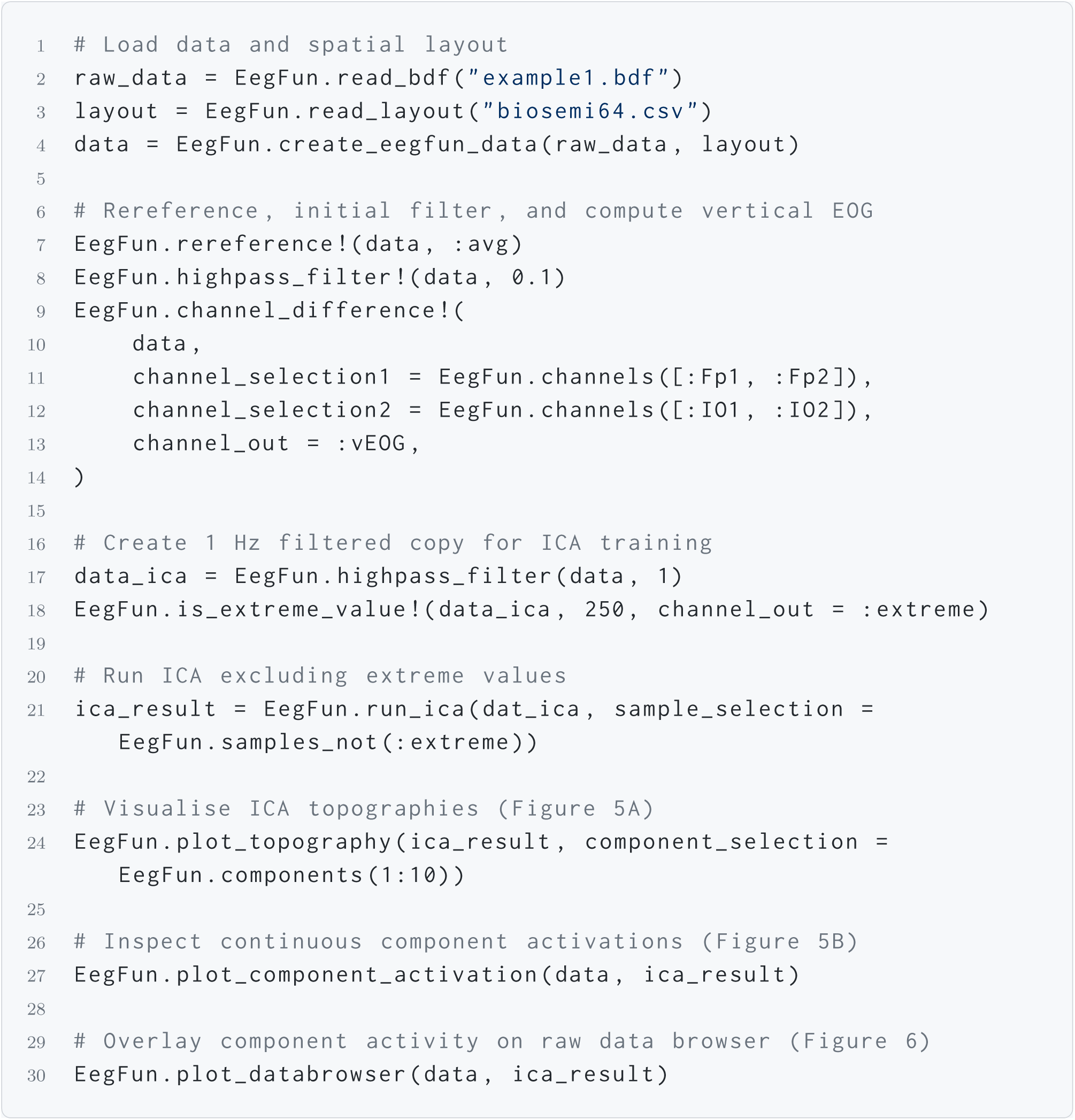
ICA pipeline demonstrating a 1.0 Hz training copy and extreme-value masking. The subsequent interactive visualisations allow researchers to inspect component spatial topographies (Figure 5A), temporal activations (Figure 5B), and the direct effect of component removal overlaid on the continuous raw EEG trace (Figure 6).

Following extraction, epoch-level baseline correction involves calculating the mean voltage across a short baseline interval (e.g., −200 ms to 0 ms) and subtracting this value from every sample in the epoch. At this stage, researchers typically employ an optional artifact detection step to identify and discard epochs containing uncorrected transients. `EegFun.jl` provides robust automated artifact detection procedures that can evaluate epochs based on absolute amplitude bounds, statistical properties (e.g., Z-scores, joint probability), or a combination of both (Listing 9).

**Listing 8:**
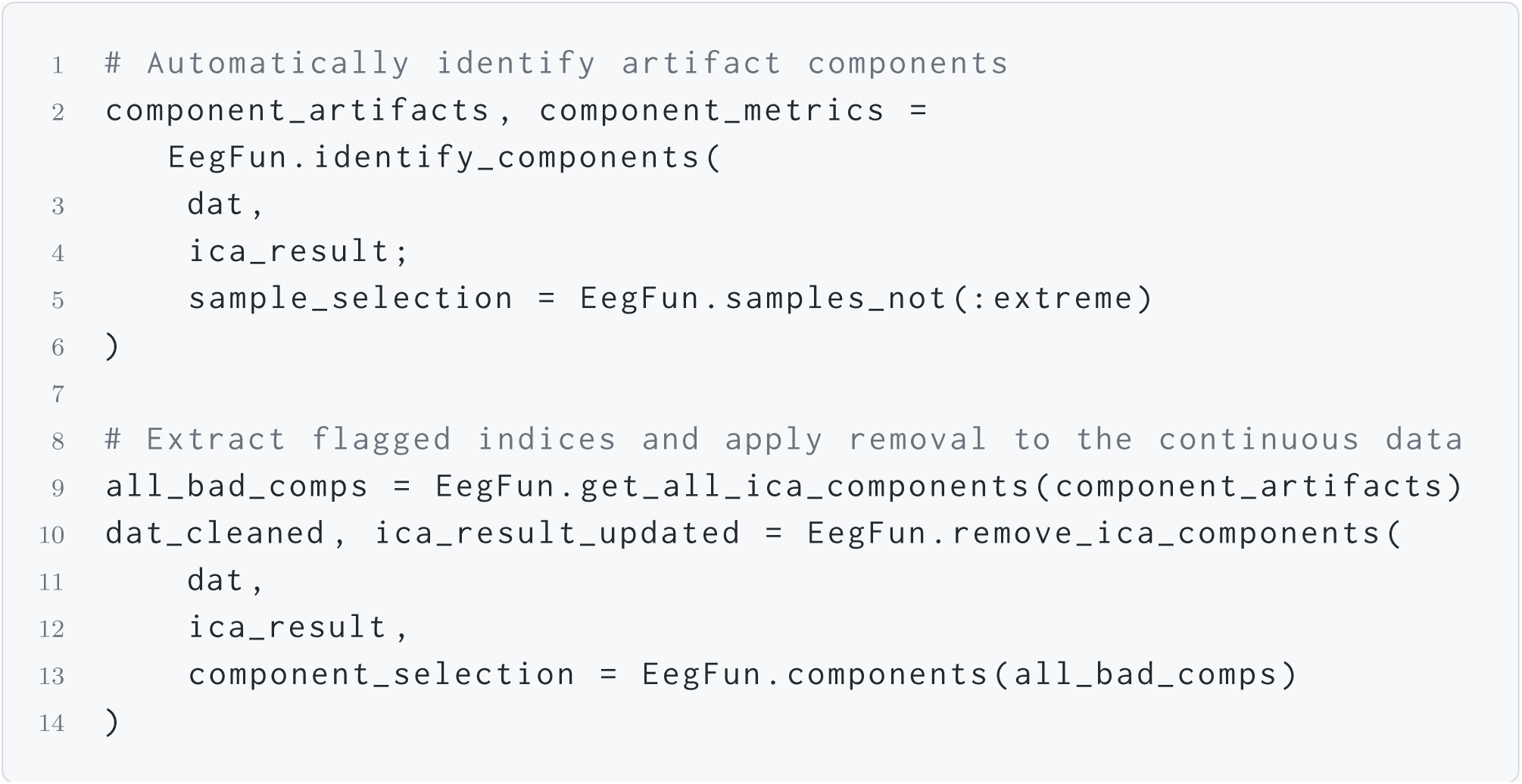
Automated identification and removal of artifactual Independent Components.

**Listing 9:**
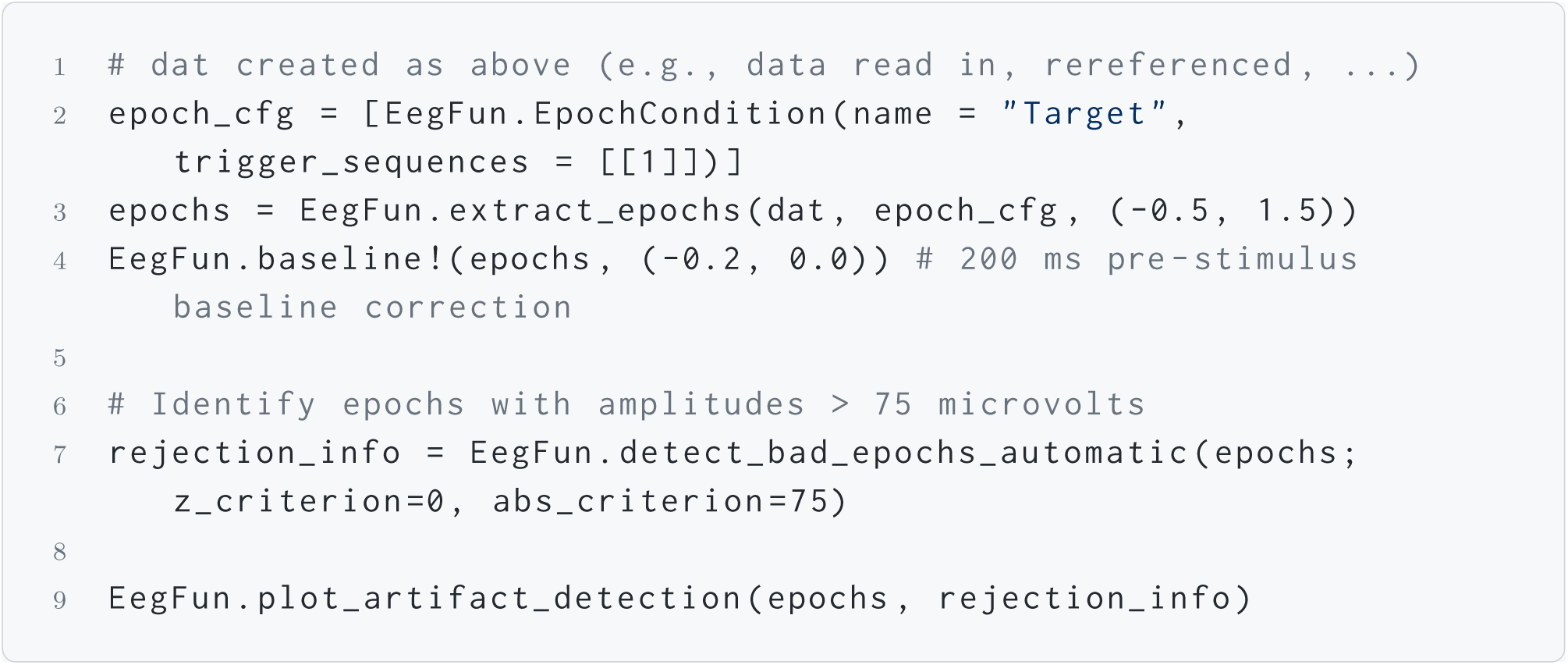
Defining extraction parameters to isolate target trials (−0.5 to 1.5 s), applying a pre-stimulus baseline correction (−200 to 0 ms), and executing automated artifact rejection to discard epochs exceeding a 75 *µ*V threshold. The final command generates an interactive visual summary (Figure 7) to inspect the artifact detection process.

A visual summary of the rejection process is shown in Figure 7, allowing the researcher to confirm that no uncorrected artifacts propagate into the final average.

**Figure 7.**
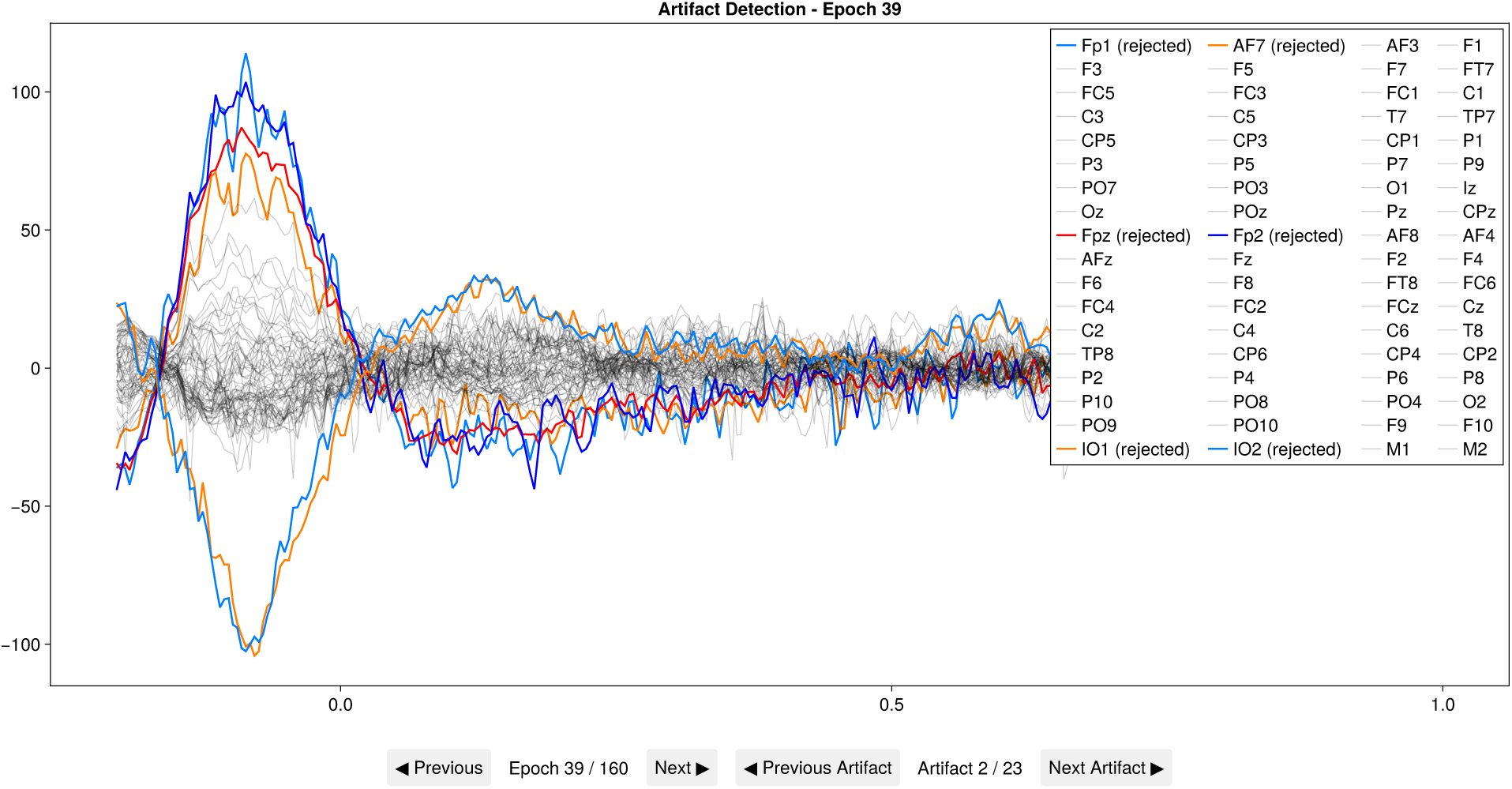
Automated artifact detection summary. Each row represents one trial; rejected epochs are highlighted, giving the researcher an audit trail of the rejection decisions.

#### Averaging and Visualisation

After trial rejection, the remaining epochs are averaged per condition and per channel to yield the final individual participant ERP waveforms. Visualising these individual-level averages is a critical quality assurance step to ensure data integrity prior to group-level aggregation. Following batch processing across an entire study cohort, these individual averages are combined to construct a grand-average ERP representing the group-level response. Visual inspection of both individual and grand-average ERPs serves as an essential sanity check to verify the presence, timing, and amplitude of expected neural components before proceeding to formal statistical inference (Listing 10, Figure 8).

**Listing 10:**
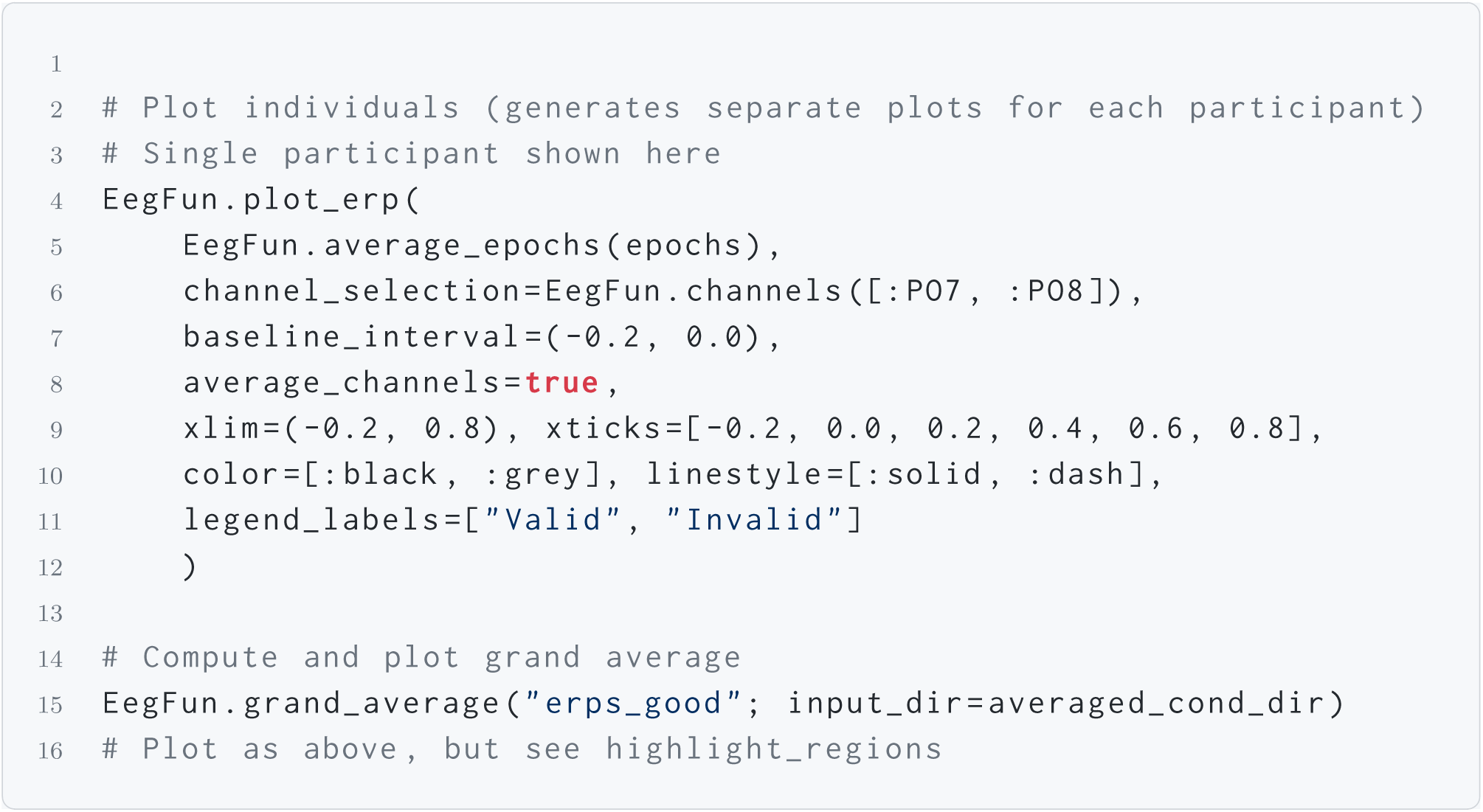
Batch filtering, computing grand averages, and generating visualisations across an entire cohort using `EegFun`’s high-level API.

**Figure 8.**
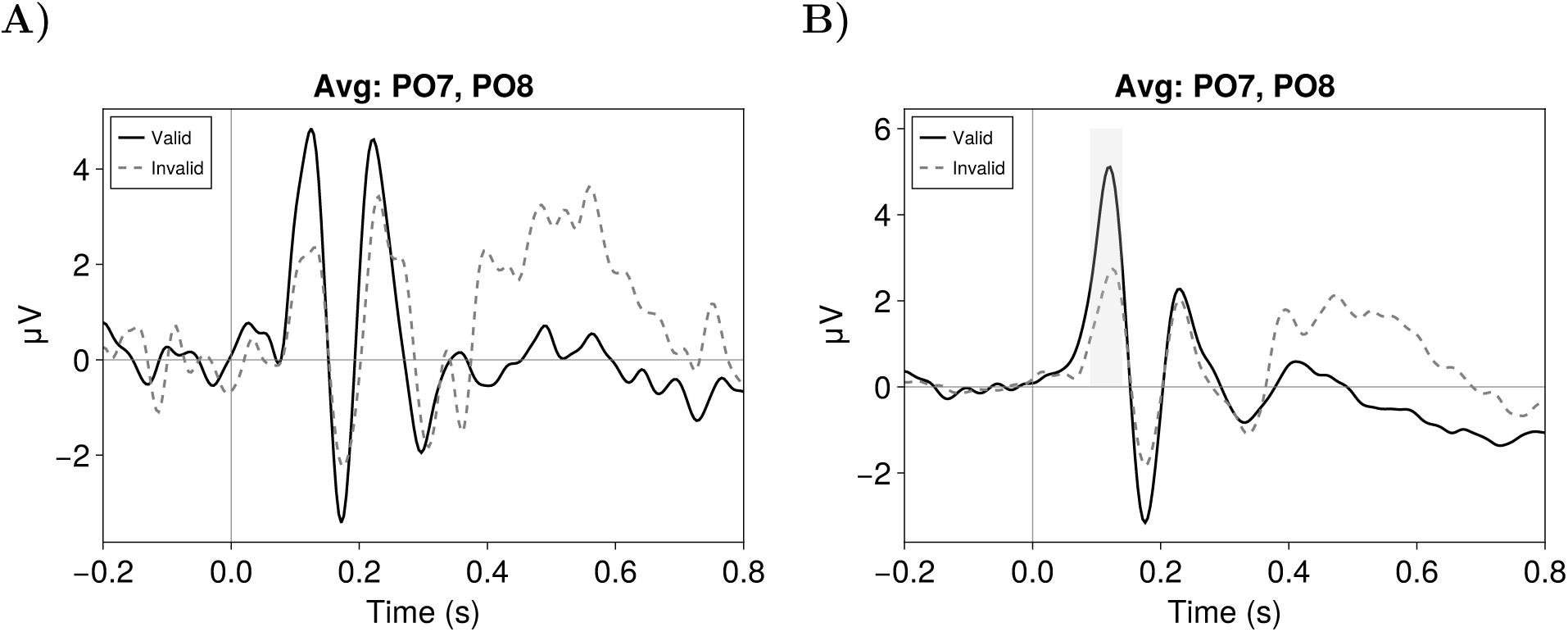
**(A)** A representative single participant’s ERP waveforms across valid and invalid conditions at averaged parieto-occipital sites (PO7/PO8). Whilst the code snippet (Listing 10) automatically generates individual plots for the entire cohort (n=12), only one is visualised here for brevity. **(B)** The resulting group-level grand-average ERP across participants. Averaging across the cohort highlights the neural components that differentiate the valid and invalid attention conditions.

#### Measurement Extraction and Export

Whilst complementary analyses such as time-frequency decomposition and multivariate decoding can be conducted natively within Julia (see the Complementary Analysis Methods section), many researchers prefer to extract discrete component measurements (such as mean amplitudes or peak latencies within a defined time window). `EegFun.jl` supports this standard workflow natively via the `erp_measurements` function. By supplying a measurement type (e.g., `“mean_amplitude“`, `“max_peak_latency“`, or `“fractional_area_latency“`), a time interval, and a channel selection, researchers can instantly quantify ERP components across all participants and conditions. Notably, the function returns a standard, “long-format” `DataFrame`. This tabular structure can be passed directly to native Julia statistical packages, such as `AnovaFun.jl` for traditional ANOVA and *t*-test comparisons, or `MixedModels.jl` (Bates et al., 2015) for linear mixed-effects modelling. Alternatively, it can be saved directly to a CSV file (Listing 11) for analysis in external software like R, JASP, or SPSS.

To aid in parameter selection and teaching, `EegFun.jl` also provides an interactive visual dashboard (`EegFun.plot_erp_measurement_gui`). This interface allows users to drag sliders to adjust the measurement and baseline windows whilst seeing the resulting calculated value (e.g., the local peak latency) dynamically updated on the plotted waveform. This visual feedback loop is particularly valuable in classroom settings to demonstrate how different baseline choices or fractional area thresholds affect the final extracted values before executing a study-wide batch extraction.

#### Data Saving (JLD2)

At any stage of the pipeline, from raw continuous structures to final ERP averages, data can be serialised to disk. `EegFun.jl` leverages `JLD2.jl`, a pure-Julia implementation of the HDF5 format, to safely write and read complex hierarchical data structures. Calling `EegFun.save_jld2(“data.jld2”, dat)` persists the entire struct, including the time-series matrix, the spatial layout, and the `AnalysisInfo` metadata history. This single-file encapsulation simplifies data management compared to multi-file formats (e.g., separate `.set` and `.fdt` files), ensuring that data and metadata remain permanently linked during sharing or archival.

**Listing 11:**
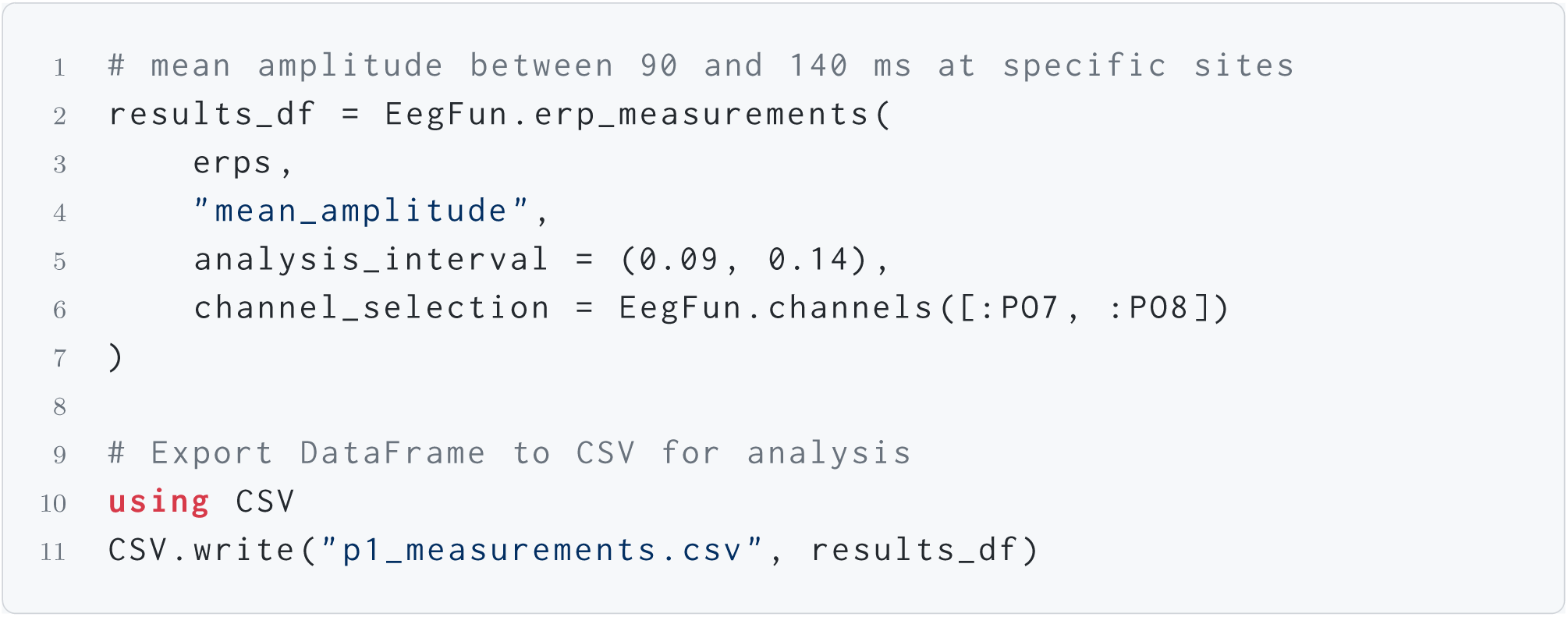
Extracting the mean amplitude of a P1 component for export.

#### High-Performance Visualisation

To maximise both interactivity and publication quality, the entire plotting suite in `EegFun.jl` is built upon `Makie.jl` (Danisch & Krumbiegel, 2021). This backend provides hardware-accelerated rendering capable of handling millions of data points without stuttering, alongside a robust declarative layout system. Researchers can export composite, multi-panel figures as resolution-independent vector graphics (e.g., SVG, PDF) or high-DPI raster images directly from the plotting functions, producing publication-ready visualisations.

### Automated Batch Pipeline

Whilst the step-by-step walkthrough illustrates the individual building blocks of the ERP workflow, in practice, multi-participant studies require processing dozens of files with identical parameter settings. `EegFun.jl` provides a fully automated batch preprocessing pipeline through the `preprocess()` function, which accepts a single TOML configuration file and iterates over all raw data files in a specified directory. The overarching processing philosophy of this pipeline aligns with the stance recently advocated by Delorme (2023), which argues that “EEG is better left alone.” Consequently, the default preprocessing strategy prioritises data integrity by avoiding aggressive, complex transformations where simpler ones suffice. However, while `EegFun.jl` encourages conservative filtering, it also recognises that artifacts must be actively managed. To this end, the pipeline provides a robust suite of tools for the rigorous detection and rejection of contaminated trials, as well as the targeted use of Independent Component Analysis (ICA) to correct for stereotypical ocular artifacts when appropriate for the experimental paradigm.

All analysis parameters, including filter cutoffs, epoch boundaries, artifact thresholds, and ICA settings, are declared once in a human-readable TOML file (Listing 12). The configuration system merges user-supplied values with a validated set of defaults, so only deviations from the defaults need to be specified. Crucially, this single configuration file acts as a complete, machine-readable provenance record for the entire analysis, ensuring 100% computational reproducibility and straightforward sharing of methods. Because TOML is a plain-text format, the configuration file integrates naturally with version control systems such as Git, allowing researchers to track every parameter change across the lifetime of a study and to include exact analysis specifications as supplementary materials alongside their publications.

This declarative configuration is complemented by an exceptionally flexible epoch extraction engine. Through the `epoch_condition_file`, researchers are not restricted to simple single-trigger extraction. Instead, they can map multiple arbitrary trigger codes to a single logical condition, or define complex multi-trigger sequences (e.g., extracting an epoch only when a specific visual stimulus is followed by a correct motor response) natively via the configuration without writing custom parsing scripts.

**Listing 12:**
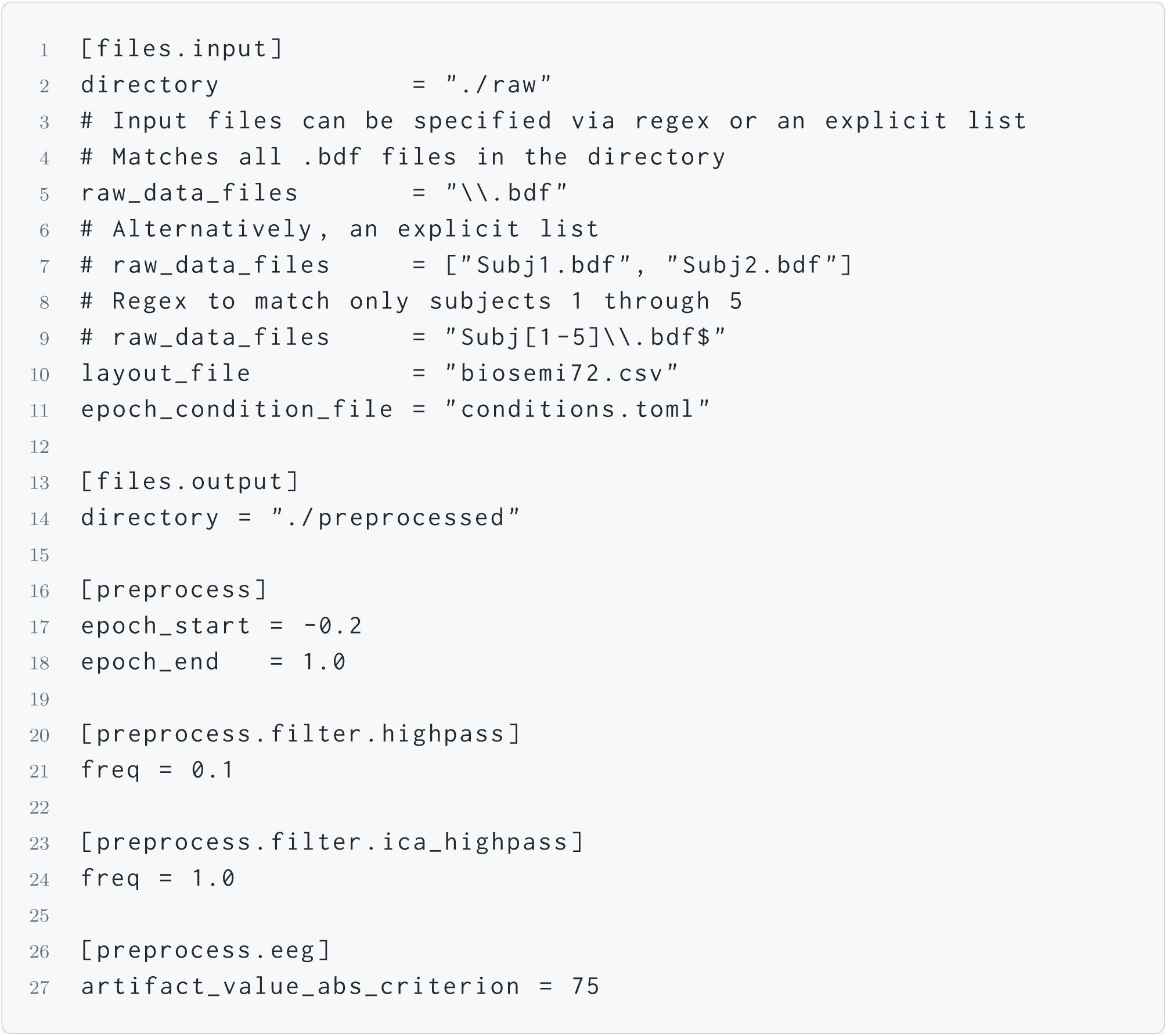
Example TOML configuration file for batch preprocessing.

**Listing 13:**
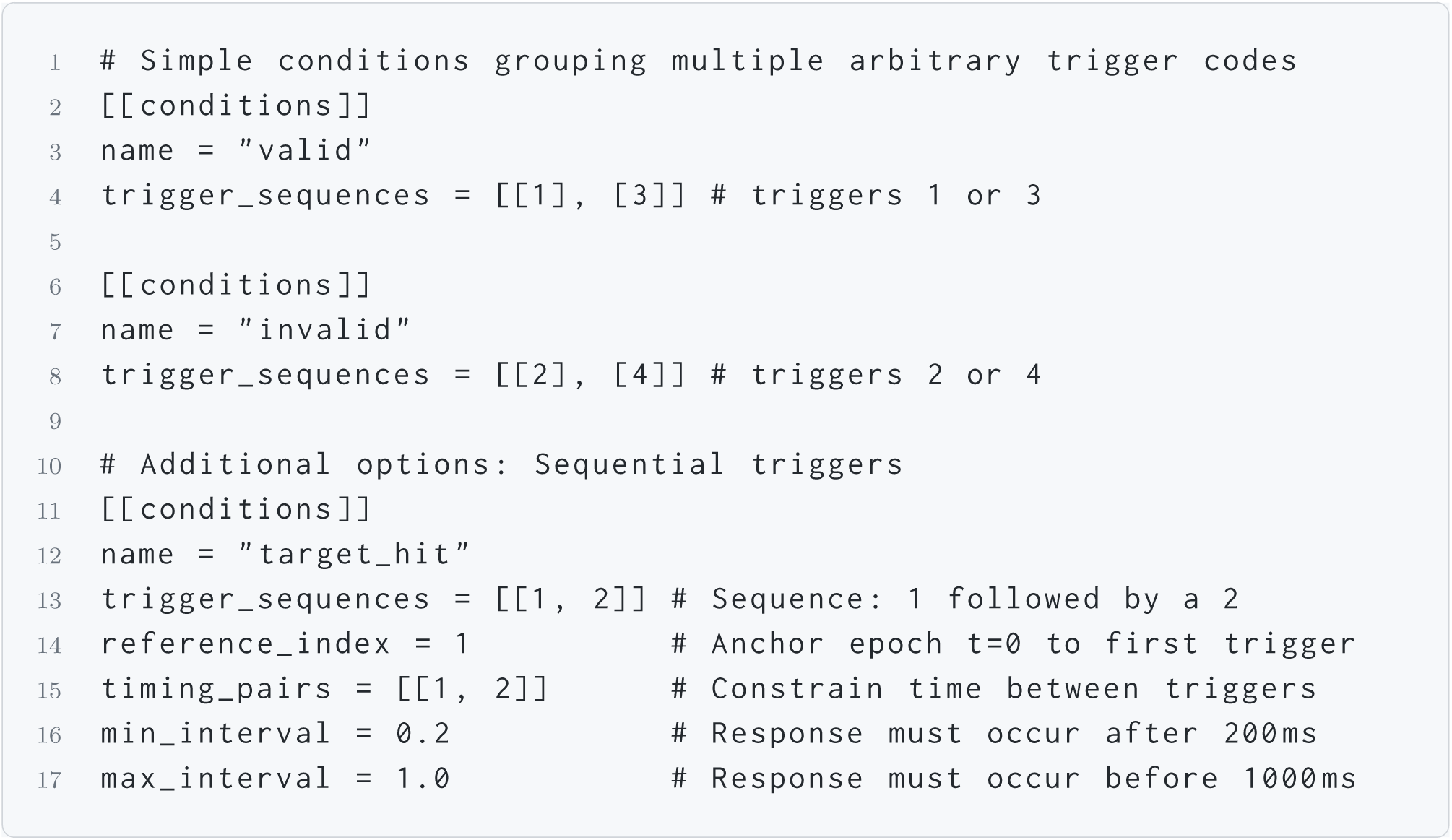
Example `conditions.toml` defining both simple grouped conditions where conditions are defined according to unique triggers, and sequential constraints including timing parameters.

With the configuration and the condition file in place, the entire multi-file pipeline is launched with a single function call (Listing 14).

The pipeline applies the complete sequence of operations described in the interactive walkthrough (see the Interactive Step-by-Step Walkthrough section), including rereferencing, filtering, ICA, bad-channel detection and repair, epoching, baseline correction, and trial rejection, to every file matched by the `raw_data_files` pattern. Intermediate and final outputs are saved to the output directory in JLD2 format at configurable stages. Depending on the configuration, these outputs include cleaned continuous data (`_continuous_cleaned`), pre- and post-rejection epochs (`_epochs_original`, `_epochs_good`), averaged ERPs (`_erps_good`), as well as the independent components (`_ica`) and comprehensive artifact tracking histories (`_artifact_info`). Additionally, the pipeline automatically generates detailed per-file and study-level summary logs. These logs provide a comprehensive audit trail of the preprocessing history, recording the exact ICA components flagged and removed (including their associated EOG correlations), a step-by-step breakdown of trials discarded during artifact rejection, and the final count of clean epochs retained per condition. Failed files are caught and reported without aborting the remaining participants, ensuring that a single corrupted recording does not halt a long overnight batch run.

**Listing 14:**
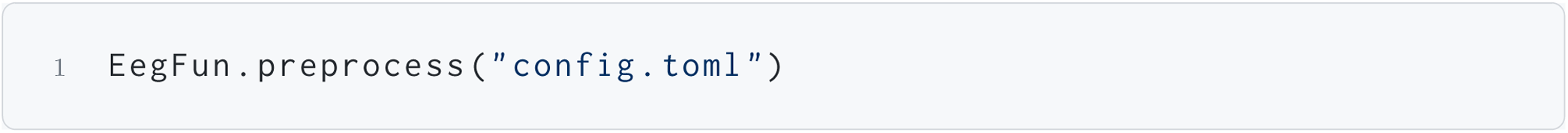
Launching the automated batch preprocessing pipeline.

For advanced users requiring logic that extends beyond the default configuration file, `EegFun.jl` includes a template generator (`EegFun.generate_pipeline_template()`). This tool produces a boilerplate Julia script containing standard setup, logging, and error handling, providing a starting point for developing customised processing workflows.

### Pipeline Outputs and Post-Processing

Once the automated batch pipeline has completed, the resulting output files are systematically organised into a designated directory structure (e.g., `preprocessed/erps_good/`). Because each file is saved in the self-contained JLD2 format, researchers can easily reload these intermediate or final datasets to perform targeted post-processing without needing to re-run the entire pipeline. For example, it is common to apply an additional low-pass filter to the final averaged ERPs to produce smoother waveforms for publication, or to compute condition-level differences across the cohort (Listing 15).

**Listing 15:**
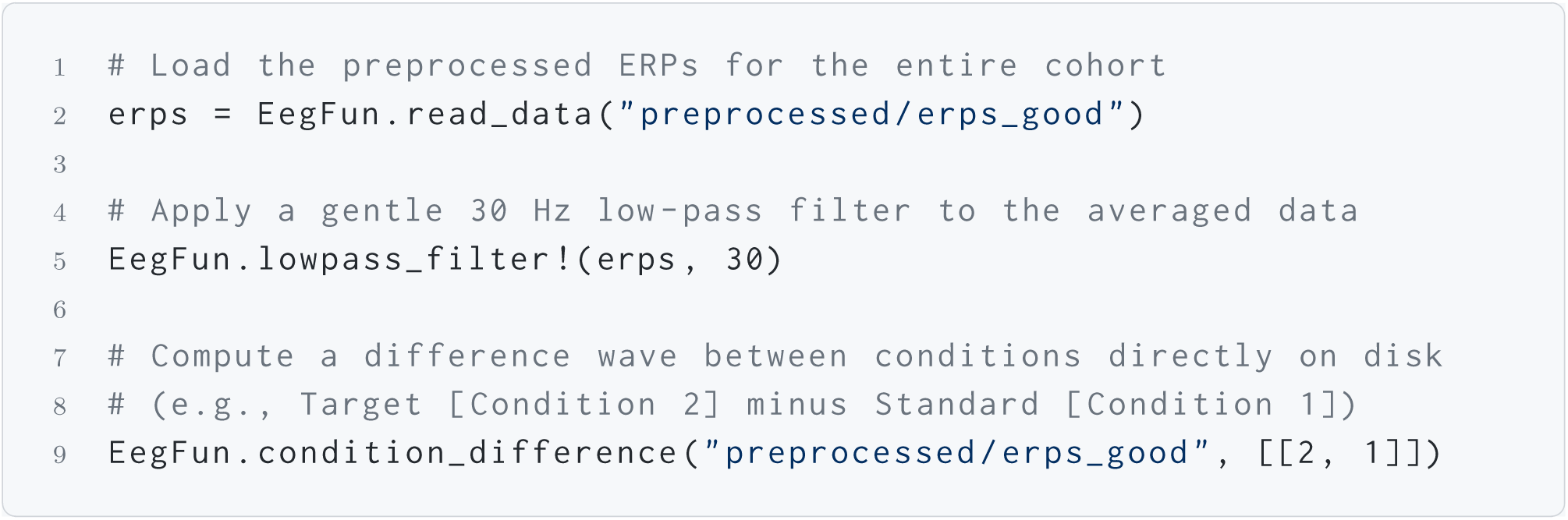
Post-processing operations applied to previously generated batch outputs. Here, final ERP files are reloaded, filtered, and subjected to condition-level arithmetic.

A fundamental principle of `EegFun.jl` is the guarantee of strict computational reproducibility and transparent provenance. Every function call that modifies the data (e.g., filtering, rereferencing, or baseline correction) is automatically recorded within the data structure’s `AnalysisInfo` metadata. When files are saved and subsequently reloaded, this exact processing history is preserved. Consequently, an analyst reviewing a specific `.jld2` file can programmatically retrieve the exhaustive list of all operations applied to that dataset from its raw origin to its current state, ensuring that the full analysis chain remains unambiguous and reproducible.

## Performance and Benchmarking

To evaluate the execution speed of EegFun.jl against existing standards, we developed a cross-pipeline benchmark comparing the time required to complete a standard preprocessing workflow (import, rereferencing, filtering, extended Infomax ICA decomposition and single-component removal, epoching, baseline correction, artifact rejection, and ERP averaging) in EegFun.jl, EEGLAB (MATLAB), and MNE-Python.

All three pipelines processed an identical dataset comprising twelve continuous `.bdf` recordings (approximately 100 MB each, sampled at 256 Hz with 72 channels) from a standard Posner spatial attention cueing paradigm (comparing ERPs elicited by targets at spatially cued “valid” versus uncued “invalid” locations). Each file was processed sequentially using comparable analysis parameters.

Benchmarks were run on a desktop workstation equipped with an AMD Ryzen 9 3900X 12-core processor, 96 GB of RAM, running Ubuntu 26.04 LTS. On this dataset, the complete pipeline completed in 919 s (*M* = 76.56, *SD* = 66.24) for `EegFun.jl` (v0.7.0; Julia 1.12.6), 2944 s (*M* = 245.36, *SD* = 274.64) for MNE-Python (v1.12.1; Python 3.14.4), and 8000 s (*M* = 665.81, *SD* = 61.76) for EEGLAB (v2026.0.0; MATLAB R2026a) (Figure 9)^3^. The resulting grand-average ERP waveforms and ICA component topographies were visually inspected and found to be highly consistent across all three toolboxes, confirming pipeline equivalence. These timings should be interpreted as indicative rather than definitive, because this relatively small sample cannot capture how each toolbox scales with massive datasets; nonetheless, they demonstrate that Julia’s in-place array operations and native compilation provide substantial speed advantages for computationally demanding preprocessing workflows. It is important to note that the scripts used for this comparison are not intended as definitive “how-to” guides for optimal ERP analysis; rather, they were designed specifically to execute a dense sequence of common processing operations to provide a rigorous, comparable workload across the three environments. The full replication code for these benchmarking pipelines, alongside the data, is available in the `benchmark/` directory of the package repository. For a complete, pedagogically focused walkthrough of this Posner attention dataset, researchers are referred to the online documentation at https://igmmgi.github.io/EegFun.jl/dev/tutorials/experiments/visual-attention.

**Figure 9.**
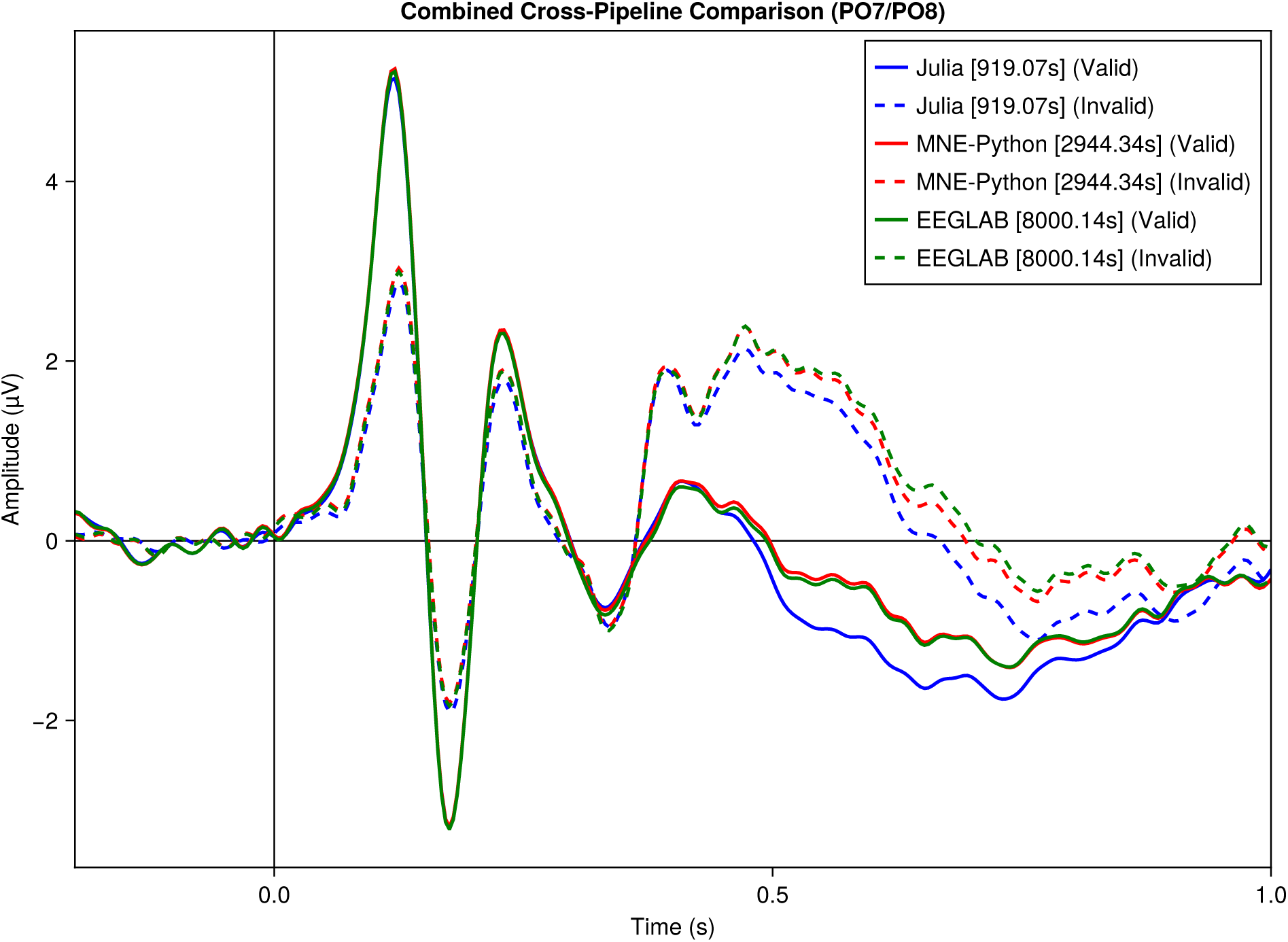
Cross-pipeline comparison of grand-average ERP waveforms (PO7/PO8) produced by `EegFun.jl` (919 s), MNE-Python (2944 s), and EEGLAB (8000 s). Execution times for the full preprocessing pipeline are shown in brackets.

## Complementary Analysis Methods

Beyond standard time-domain averaging, EegFun.jl provides integrated implementations for spectral and multivariate analysis workflows.

### Time-Frequency Decomposition

Oscillatory brain dynamics are not strictly phase-locked to a stimulus and are often cancelled out in a standard ERP average (Cohen, 2014). `EegFun.jl` provides three methods to recover this time-varying spectral content: Short-Time Fourier Transform (STFT), Multitaper estimation, and Morlet wavelet convolution. Whilst the package supports advanced configurations such as adaptive cycle bounds to optimise time-frequency resolution trade-offs, Listing 16 and Figure 10 demonstrate a fixed 3-cycle Morlet decomposition (replicating Figure 13.14 A from Cohen (2014)). Because raw spectral power inherently follows a 1*/f* distribution, `EegFun.jl` provides a dedicated `tf_baseline` function offering multiple normalisation methods (including decibel conversion, percentage change, and Z-scoring) to prepare the data for statistical inference.

**Figure 10.**
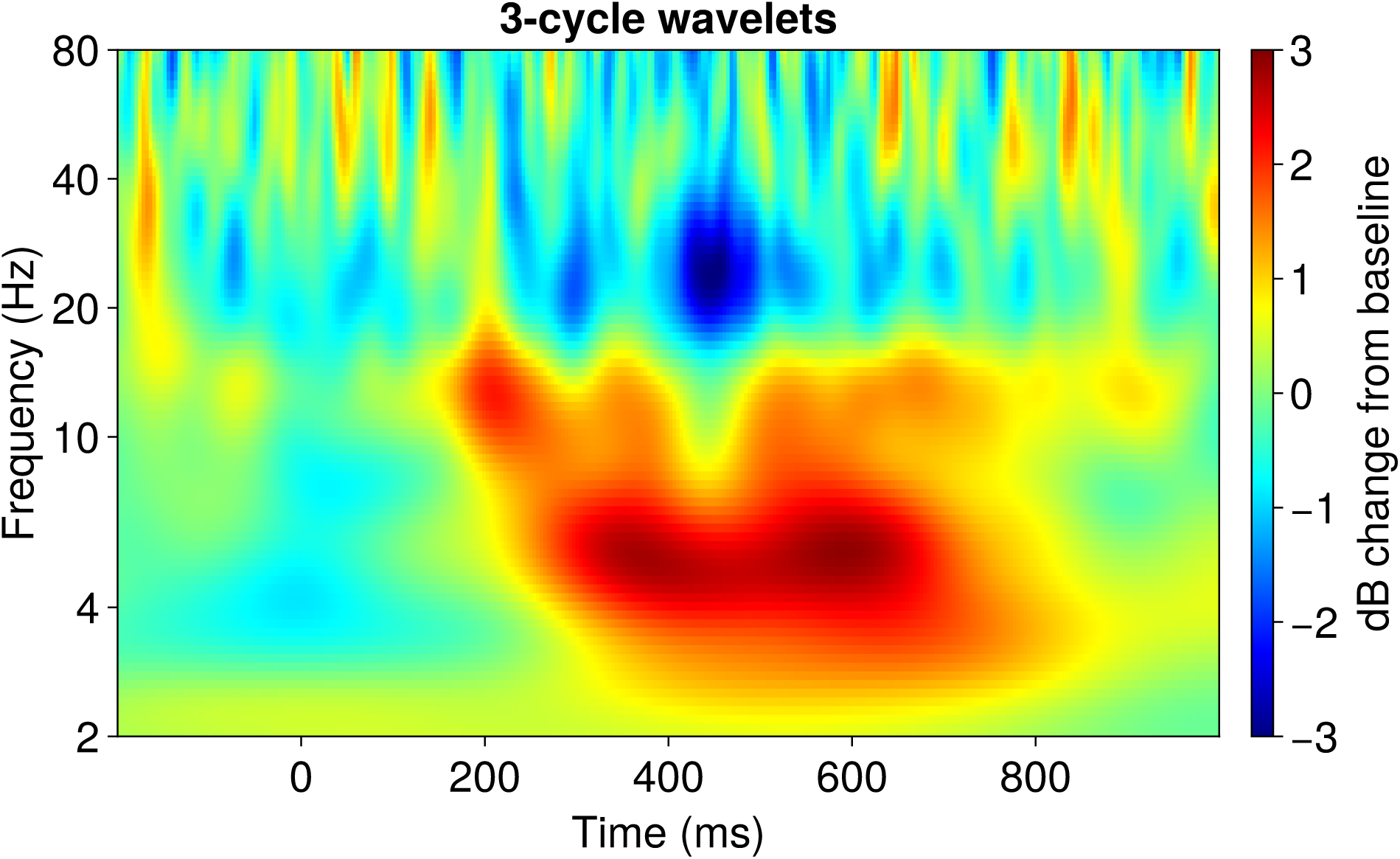
Time-Frequency power spectrum computed via Morlet wavelet convolution, with decibel baseline normalisation applied. The data, parameters, and plot settings are chosen to reproduce Figure 13.14 Panel A from Cohen (2014) (see Listing 16).

### Multivariate Pattern Analysis (MVPA)

Mass-univariate statistics test each channel independently, which limits their sensitivity to distributed neural representations. In contrast, time-resolved MVPA leverages the entire scalp topography simultaneously, applying a machine-learning classifier at every time sample to determine if the spatial pattern of voltage reliably categorises the trial condition. As demonstrated in foundational work, such multivariate approaches are highly sensitive because they bypass the need to select *a priori* channels of interest and can capture subtle, distributed signals across the scalp to track continuous cognitive processes, such as the maintenance of working memory representations or shifts in spatial attention (Bae & Luck, 2018). `EegFun.jl` leverages `LIBSVM.jl`—a Julia interface to the highly-optimised LIBSVM library (Chang & Lin, 2011)—to provide a decoding interface that handles cross-validation, iteration, and cluster-based statistical inference automatically (Listing 17, Figure 11).

**Figure 11.**
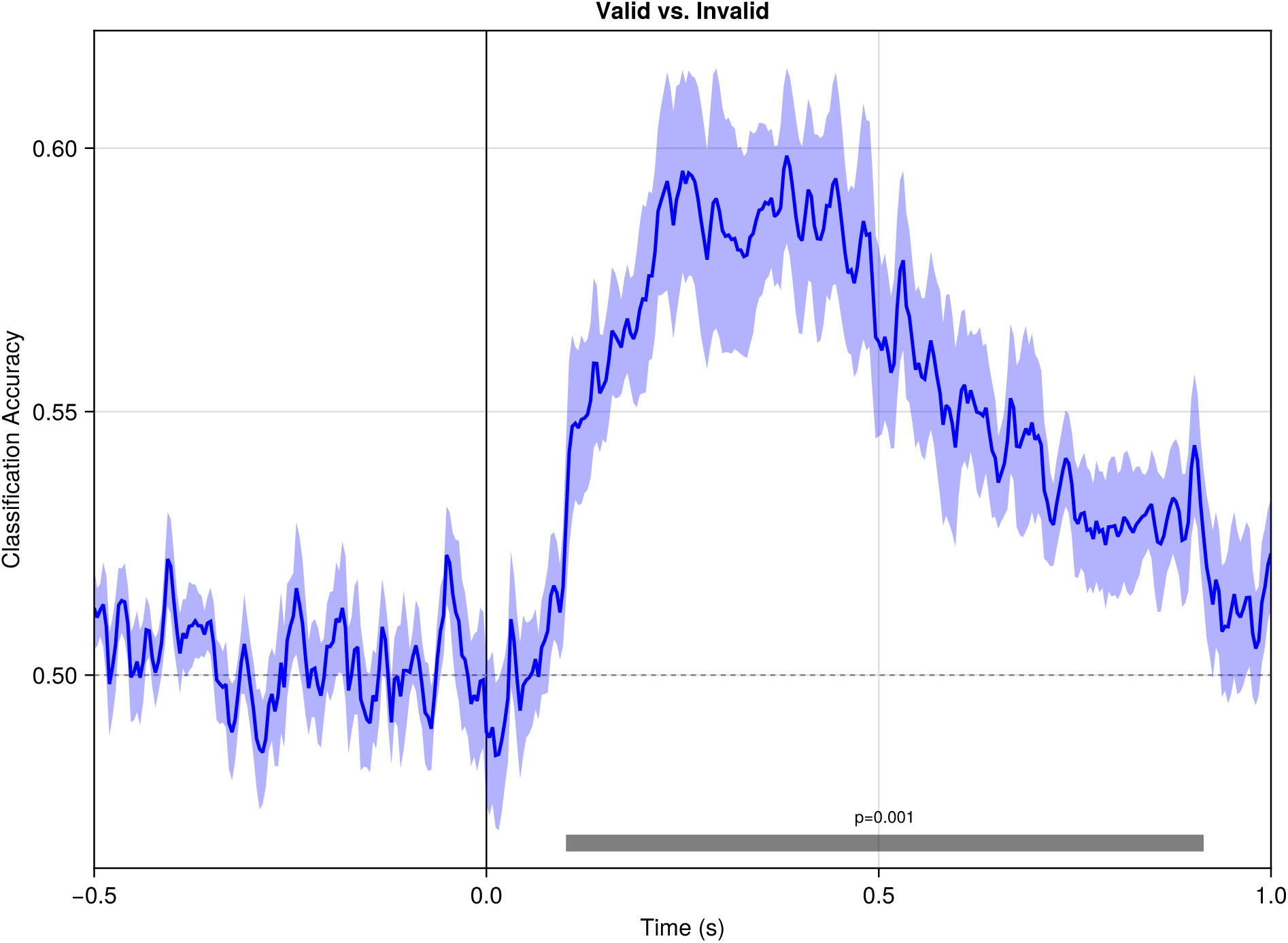
Time-resolved MVPA decoding accuracy across participants. Shaded regions indicate temporal clusters where classification accuracy significantly exceeded the 50% chance level (p < 0.05, cluster-based permutation test).

**Listing 16:**
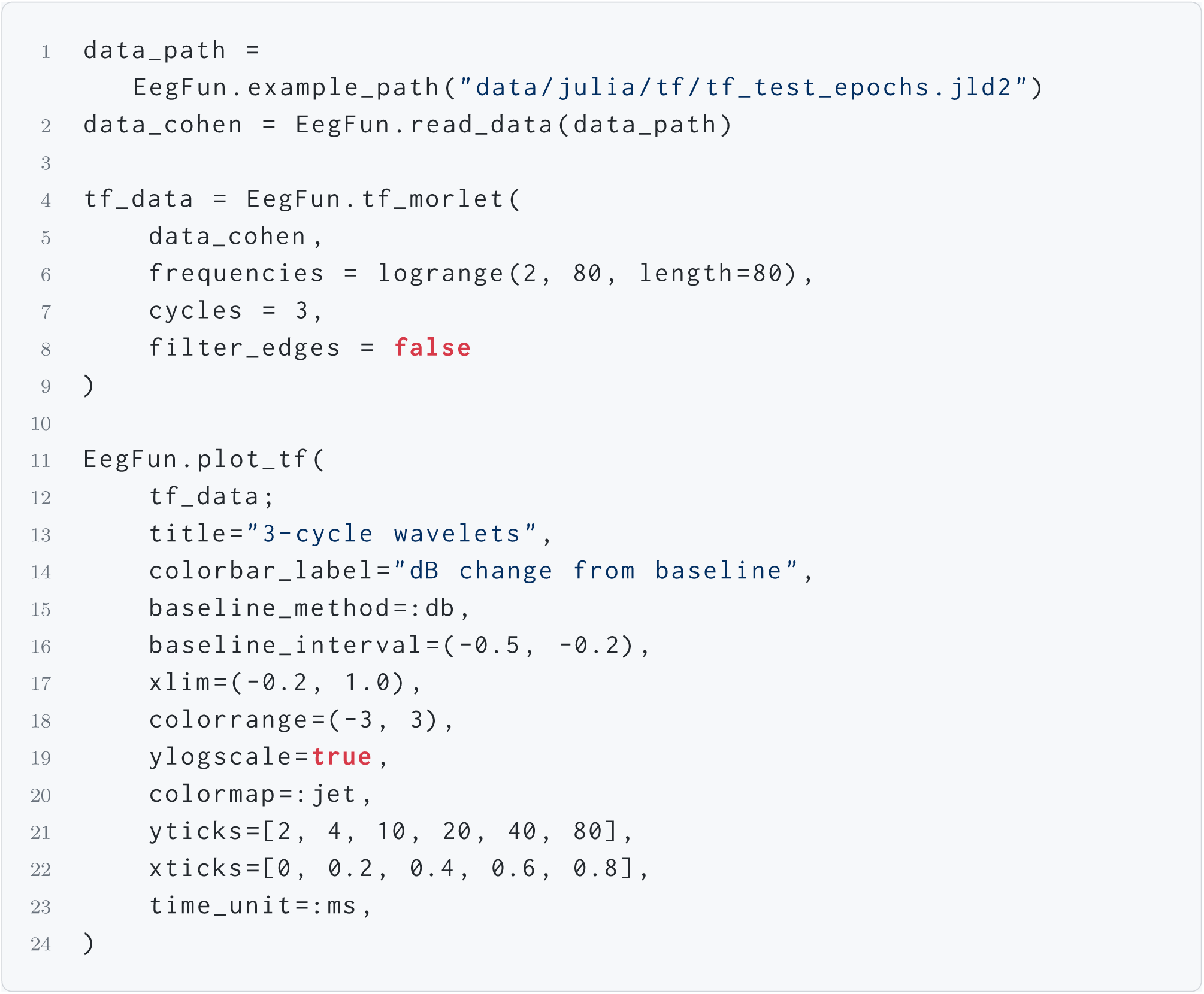
Time-frequency decomposition using 3-cycle Morlet wavelets with decibel baseline normalisation (Figure 10). The example data is from Cohen (2014), with parameters and plot settings chosen to reproduce Figure 13.14 Panel A from the same text.

**Listing 17:**
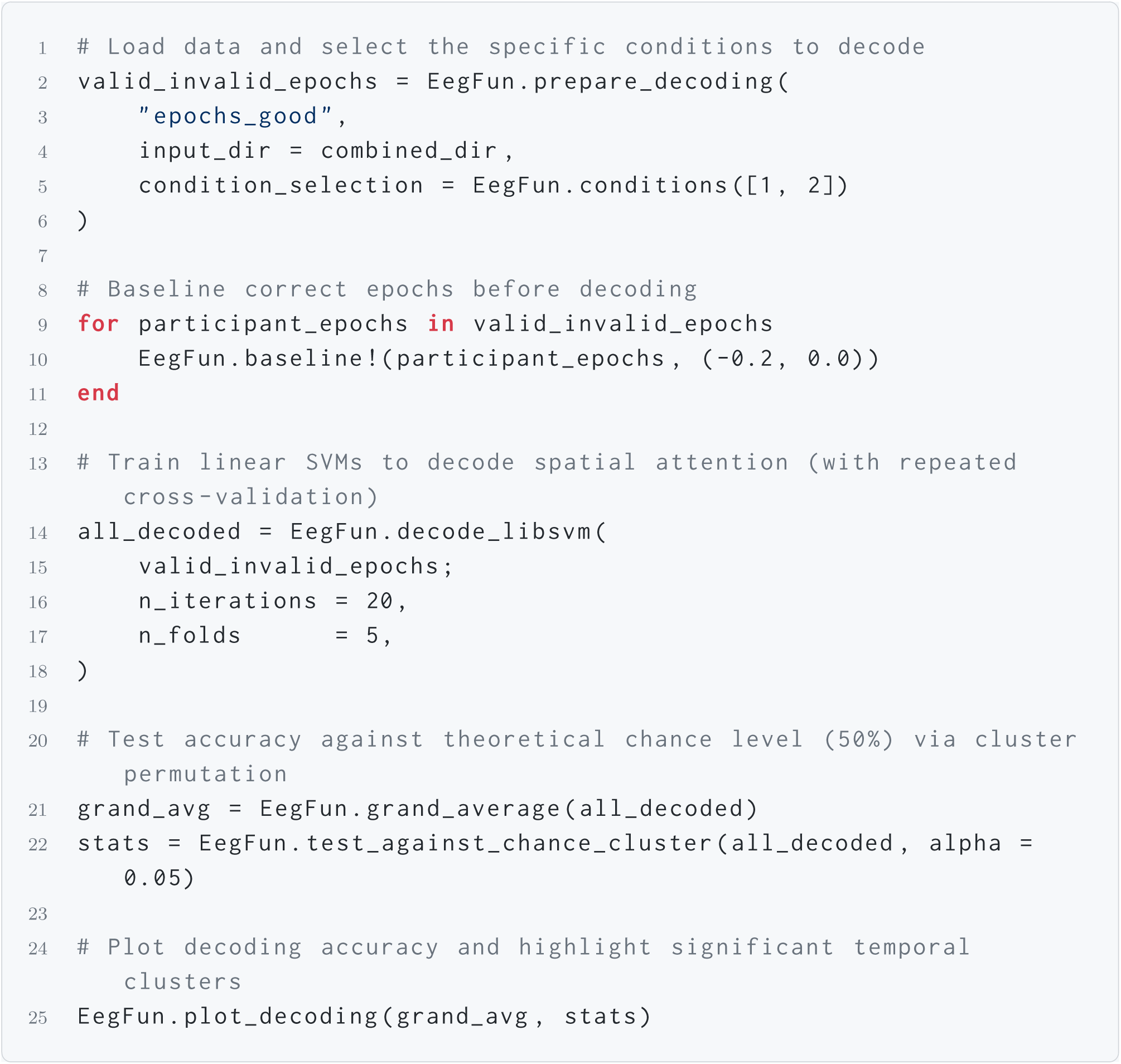
Time-resolved MVPA decoding of spatial attention (Valid vs. Invalid) across multiple participants.

## Discussion

`EegFun.jl` provides researchers with a complete, open-source package for EEG and ERP analysis implemented entirely in a single, high-level language. The package covers the full sensor-space analysis pipeline: data import from seven standard file formats (BDF, EDF, XDF, FIF, EEGLAB, FieldTrip, BrainVision), signal conditioning through zero-phase FIR filtering and rereferencing, artifact management via Infomax ICA with automatic EOG component identification, flexible epoch extraction with configurable trigger-sequence matching, baseline correction, and threshold-based trial rejection with ERP averaging. Beyond the core time-domain pipeline, `EegFun.jl` extends into time-frequency decomposition via Morlet wavelet, Short-Time Fourier Transform, and multitaper methods, as well as multivariate pattern analysis (MVPA) with cross-validated SVM decoding and cluster-based statistical inference. The entire visualisation suite—including an interactive continuous data browser, ERP waveform and topographic plots, ICA component inspection, ERP image heatmaps, and time-frequency spectrograms—is built on `Makie.jl`, providing both GPU-accelerated interactivity and publication-quality vector-graphics export.

Several design features distinguish `EegFun.jl` from existing platforms. The enforced coupling of time-series data with spatial channel layouts at construction time prevents the silent metadata loss that can occur when data and layout are held as separate variables—a common source of difficult-to-diagnose errors in workflows where data reordering or channel subsetting can silently break the correspondence between signal columns and channel positions. `EegFun.jl`’s typed data structures make such mismatches an early run-time failure rather than a silent numerical artifact in the final result. The predicate-based selection API (see the Data Selection and Predicates section) provides a composable, self-documenting subsetting mechanism that replaces fragile numeric indexing with symbolic channel and sample references that remain valid regardless of data reordering or layout changes. The TOML-driven batch pipeline (see the Automated Batch Pipeline section) offers a declarative, version-controllable specification of every preprocessing decision, enabling full computational reproducibility from a single configuration file that can be shared directly as supplementary material. Finally, by exploiting Julia’s multiple dispatch system, the same function names (e.g., `baseline`, `highpass_filter`, `plot_erp`) operate correctly across continuous, epoched, and averaged data structures, keeping the API surface small and the learning curve shallow.

`EegFun.jl` joins a small but growing ecosystem of Julia-based cognitive neuroscience tools, such as the `Unfold.jl` package for regression-based deconvolution (Ehinger & Dimigen, 2019), and aims to complement rather than replace existing analysis platforms. Being implemented natively in Julia provides a fully open-source, license-free alternative to MATLAB-based tools like EEGLAB (Delorme & Makeig, 2004) and FieldTrip (Oostenveld et al., 2011). Compared to MNE-Python (Gramfort et al., 2013), `EegFun.jl` offers an architecturally unified alternative in which both the user-facing API and the underlying numerical algorithms are expressed in the same language, simplifying debugging and lowering the barrier to methodological contributions.

Beyond research applications, `EegFun.jl` is designed with pedagogical utility in mind. The package includes a suite of interactive, `Makie.jl`-based teaching demonstrations covering core digital signal processing concepts such as Nyquist sampling, frequency-domain convolution, and spectral leakage. These visual tools allow instructors to demonstrate the mathematical foundations of EEG analysis in real-time, bridging the gap between theoretical principles and applied neuroscience. Combined with the interactive data browser, intuitive plotting functions, and the extensive online documentation featuring worked experiment walkthroughs, `EegFun.jl` is well-suited for both active research laboratories and classroom instruction.

Whilst `EegFun.jl` provides a comprehensive pipeline for sensor-space ERP and time-frequency analyses, it is important to acknowledge areas where the package currently trails behind more mature ecosystems. At present, `EegFun.jl` does not support source-space reconstruction (forward/inverse modelling), connectivity measures (e.g., phase-locking value, coherence), real-time data streaming for brain–computer interfaces, or integration with other neuroimaging modalities such as MEG or fMRI. The package is deliberately scoped to focus on more standard EEG/ERP analyses. Whilst the package’s architecture is designed to accommodate such extensions and future development may explore some of these areas, as an open-source project, community contributions to expand these capabilities are highly encouraged via the public GitHub repository. Furthermore, new users should be aware of Julia’s just-in-time compilation model. Whilst the final execution is highly optimised, the compilation overhead on the very first execution of a session (commonly referred to as the “time-to-first-X” or TTFX) can initially make interactive data exploration or plotting feel sluggish compared to interpreted languages, though this latency vanishes entirely on all subsequent calls.

## Conclusion

We have presented `EegFun.jl`, an open-source Julia package that provides a complete, scripted EEG/ERP analysis pipeline. By implementing all components, from file reading to statistical inference, in pure Julia, the package balances accessibility and performance. The consistent, typed data-structure hierarchy and the enforced coupling of time-series data with spatial layouts reduce the opportunity for silent preprocessing errors and improve the reliability of the analysis chain. We hope that `EegFun.jl` lowers the barrier to fully reproducible, open-source EEG research within the cognitive neuroscience community.

## Data and Code Availability

The source code for `EegFun.jl` is available under the MIT License on GitHub at https://github.com/igmmgi/EegFun.jl. The datasets and layouts used for the tutorials and benchmarks in this paper are included within the package and can be accessed using the `EegFun.example_path()` utility function. Additional, raw example data is hosted on Zenodo at https://zenodo.org/records/21770157. All documentation, including full code examples, is available at https://igmmgi.github.io/EegFun.jl/.

## Acknowledgements

Carolin Dudschig was funded by the German Research Foundation (DFG), Heisenberg grant (Project number: 419439493) and supported by the equal opportunity funds, DFG Research Unit 2718 (Project number: 381713393).

## Footnotes

1 Example datasets and layouts are available within the package and can be accessed via example_path raw_bdf = EegFun.read_bdf(EegFun.example_path(“data/bdf/example1.bdf”)) layout = EegFun.read_layout(EegFun.example_path(“layouts/biosemi/biosemi72.csv”)).

2 A more extreme high-pass filter (e.g., 1.0 Hz) is often required for ICA decomposition. However, the spatial weights calculated from this more aggressively filtered data are subsequently applied back to the data with more standard EEG high-pass filter settings.

3 The bottleneck in the reported benchmarks here is the ICA (in this case, extended Infomax). Note that users with a modern GPU can set the `use_gpu` flag to `true` for significant performance improvements across the currently available ICA algorithms. For example, re-running the Julia pipeline with `use_gpu = true` and the Picard ICA algorithm reduces the processing time to 40 seconds for the whole analysis run.

